# Local thermodynamics governs the formation and dissolution of protein condensates in living cells

**DOI:** 10.1101/2021.02.11.430794

**Authors:** Anatol W. Fritsch, Andrés F. Diaz-Delgadillo, Omar Adame-Arana, Carsten Hoege, Matthäus Mittasch, Moritz Kreysing, Mark Leaver, Anthony A. Hyman, Frank Jülicher, Christoph A. Weber

## Abstract

Membraneless compartments, also known as condensates, provide chemically distinct environments and thus spatially organize the cell. A well-studied example of condensates is P granules in the roundworm *C. elegans* which play an important role in the development of the germline. P granules are RNA-rich protein condensates that share the key properties of liquid droplets such as a spherical shape, the ability to fuse, and fast diffusion of their molecular components. An outstanding question is to what extent phase separation at thermodynamic equilibrium is appropriate to describe the formation of condensates in an active cellular environment. To address this question, we investigate the response of P granule condensates in living cells to temperature changes. We observe that P granules dissolve upon increasing the temperature and recondense upon lowering the temperature in a reversible manner. Strikingly, this temperature response can be captured by *in vivo* phase diagrams which are well described by a Flory-Huggins model at thermodynamic equilibrium. This finding is surprising due to active processes in a living cell. To address the impact of such active processes on intra-cellular phase separation, we discuss temperature heterogeneities. We show that, for typical estimates of the density of active processes, temperature represents a well-defined variable and that mesoscopic volume elements are at local thermodynamic equilibrium. Our findings provide strong evidence that P granule assembly and disassembly are governed by phase separation based on local thermal equilibria where the non-equilibrium nature of the cytoplasm is manifested on larger scales.

**SIGNIFICANCE STATEMENT:** Living cells rely on a continuous flux of energy to spatially organize biochemical processes. It remained unclear whether cells can achieve this spatial organization via thermodynamic principles. Here, we report the striking behavior of a cold-blooded organism that reacts to environmental temperature changes similar to a thermodynamic system at local equilibrium. Our key finding is that protein-rich droplets form and dissolve reversibly with temperature due to changes in the organism?s entropy. We show that the organism uses a specific molecule to extend droplet stability to the natural temperature range of the organism’s habitat. Due to the relevance of such protein droplets for the organism?s fertility, our works shed light on how molecular components could facilitate biological functions via thermodynamic principles.

## I. INTRODUCTION

Living cells use metabolic fuel in the form of ATP or GTP to express proteins, run enzymatic reactions and maintain the activity of chaperones and molecular motors. These fuel-driven, metabolic processes keep cells away from thermodynamic equilibrium and thereby allow the assembly of functional assemblies and structures that are not possible at thermodynamic equilibrium. Despite the relevance of metabolic processes to the physiology of living cells, evidence is accumulating that the physics of phase separation provides a good description for the formation of many intracellular condensates. Potential candidates are protein-RNA condensates, such as P granules and stress granules, which assemble via phase separation in the cyto- or nucleoplasm in various different cell types [1–5]. A decade ago, a seminal study in cell biology has revealed that these condensates share the physical properties of phase-separated, liquid-like droplets [6]. In a typical phase separation picture, different phases coexist at thermodynamic equilibrium. However, a cell uses a considerable amount of ATP and GTP which is hydrolyzed to fuel enzymatic reactions. A particular ex-ample are DEAD-box ATPases [7], which were shown to promote phase separation in their ATP-bound form, whereas ATP hydrolysis induces compartment turnover and release of RNA [8]. These metabolic processes raise the question of whether it is appropriate to use equilibrium thermodynamics to discuss the formation of condensates in living cells. Such questions also arise for individual molecules. For instance, molecular motors use ATP to move along cytoskeletal filaments [9]. However, such motors operate isothermally because temperature fluctuations relax within nanoseconds at the nanometer scale of macromolecules [10]. Thermodynamic phase coexistence requires at least the phase boundary to be locally at equilibrium [11]. A key question is whether such conditions are satisfied in a cell given the continuous consumption of fuel molecules such as ATP or GTP. Such questions can be tested by examining the response of condensates using temperature as a control variable for phase separation. Interestingly, qualitative experimental studies in the *C. elegans* roundworm suggest that P granule condensates can dissolve upon increasing the temperature and recondense when the temperature is lowered again [12, 13]. Here, we investigate the question of whether the temperature response of protein-RNA condensates can be understood by the thermodynamics of phase separation. We study P granule condensates in embryonic cells of the roundworm *C. elegans*, which is a cold-blooded organism that adapts and responds to the temperature of the environment [14] (Fig. 1A). We subject early one-cell embryos to temperature changes controlled by a thermostat and quantitatively analyze the dynamics of P granules. We show that this system can be described by a thermodynamic Flory-Huggins model for phase separation.

**FIG. 1.**
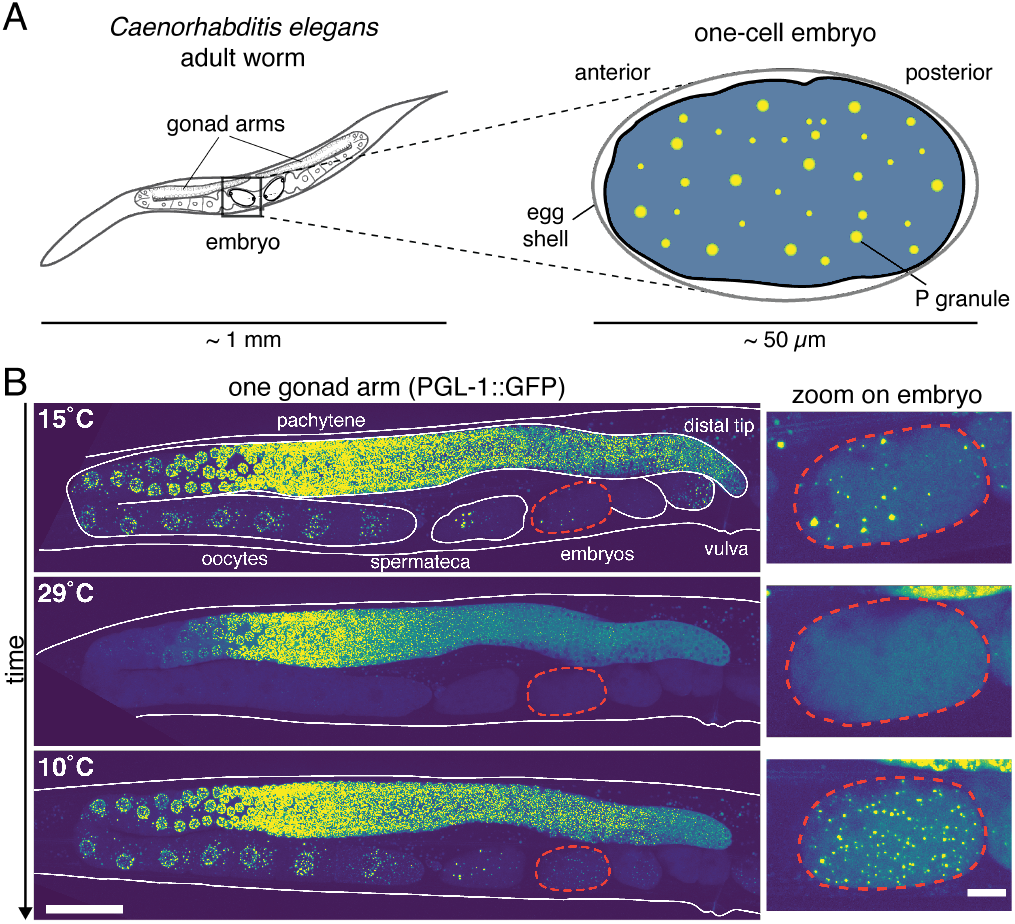
Illustration of a *C. elegans* roundworm and reversible P granule dissolution upon heating. A. We consider the response of *µm*-sized P granules in the one-cell stage of the *C. elegans* embryo against temperature changes to exemplify the role of thermodynamics for the formation and dissolution of protein condensates in a living organism. B. P granules labeled via PGL-1::GFP (yellow-green dots) dissolve upon heating and reform when cooled again within a few minutes both in embryos as well as in parts of the gonad (scale bars 50 *µm* (left) and 10 *µm* (right)).

## II. RESULTS

### A. Reversible condensation and dissolution upon temperature changes

Figure 1B shows the response of P granules (labelled via PGL-1::GFP which we refer to as wildtype) in a *C. elegans* worm to a temperature shift from 15 to 29 and then down to 10 degrees Celsius. P granules dissolve upon temperature increase and reform when cooled, in agreement with previous observations [12, 13]. In addition, we see a similar behaviour in parts of the gonad (distal tip, oocytes), while the pachytene region seems more robust against the transient temperature change. In the following experiments, we dissect single embryos from the worm for better imaging and mount them in a chamber with temperature controlled by a thermostat (SI, Fig. S1). To quantify the temperature response, we subjected embryos to repeated cycles of temperature increase and decrease (Fig. 2A). We were able to observe up to 5 cycles of dissolution and condensation showing that the formation of P granules in the *C. elegans* embryo is reversible over multiple cycles (Fig. 2A; SI movie S1, description SI sect S7). Together with previous experiments, this suggests that P granules form and dissolve by thermodynamic phase separation.

**FIG. 2.**
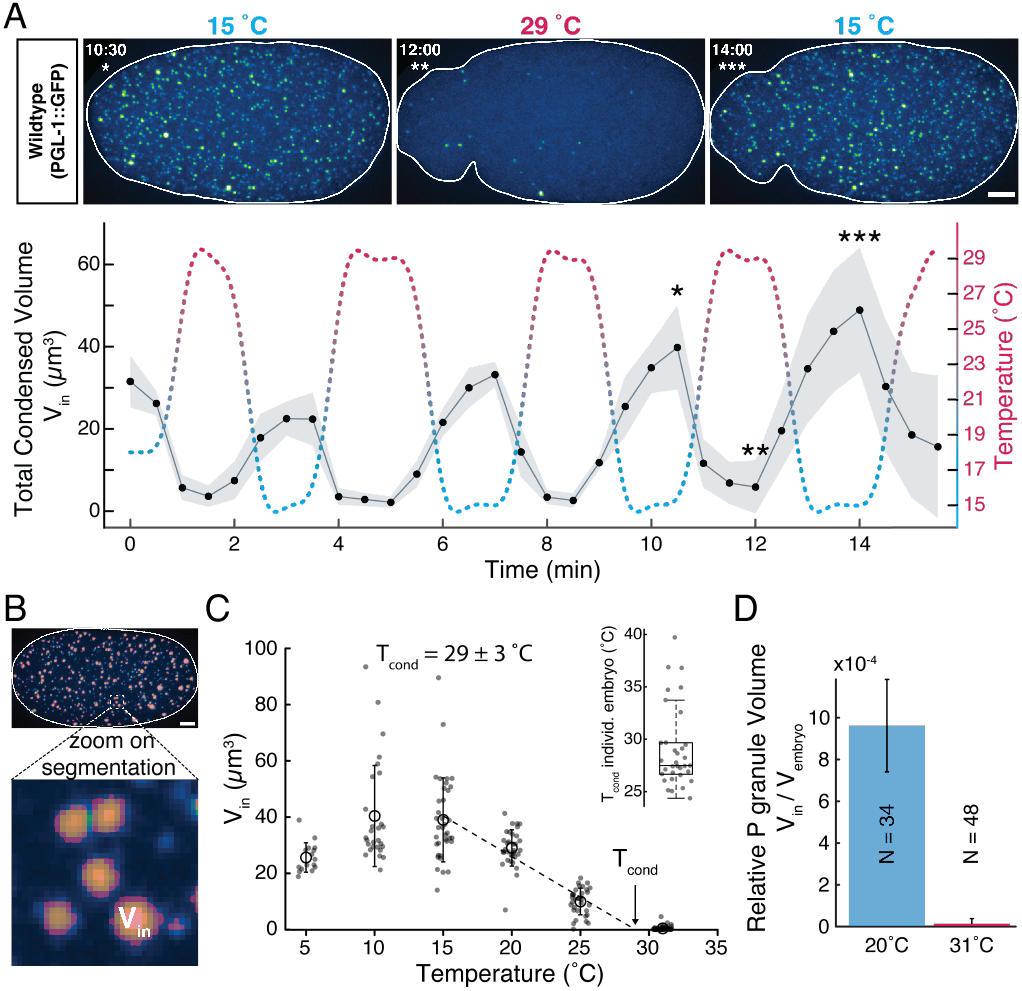
P granule volumes respond to temperature oscillations and determine the global condensation temperature. A. Five oscillations in temperature from 15 to 29 °C cause an in-phase oscillation of the total condensed volume of P granules, *V*_in_ (see example embryo going through one cycle, n=4, shaded area is STD, scale bar: 5 *µm*). The notable increase of total condensed volume with the number of cycles is likely due to the progress of one-cell embryos from an early homogeneous to a later polarized state. B. Mask used to extract the P granule volumes from the microscope images (details on image analysis, see materials and methods and SI). C. The condensation temperature *T*_cond_ is defined as the temperature where the total volume *V*_in_ averaged over all embryos becomes zero. Inset: We obtain a very similar value for condensation temperature when extracting *T*_cond_ for each embryo and averaging over all considered embryos. D. At *T* = 20 °C condensates contribute to approximately 1 · 10^*-*3^ of the volume of an embryo (*N* number of embryos per condition, errorbars are SD). Around the dissolution temperature, the relative volume of P granule phase drops to values close to zero.

The response of a phase separating system to temperature provides important information about a number of thermodynamic parameters. One important parameter is the temperature at which the system mixes/demixes, which we call the condensation temperature *T*_cond_. This is the temperature at which the average concentration of PGL-1 equals the saturation concentration. Above this temperature the system stays under-saturated and no P granules form. Below this temperature the system is super-saturated and P granules form. As one approaches the condensation temperature from below, the total volume of P granules should decrease in a linear manner toward zero.

We determine the total P granule volume *V*_*in*_ using a segmentation mask, see Fig. 2B and materials and methods. Fig. 2C shows a linear decrease of *V*_*in*_ between 15°C and 30°C, implying a condensation temperature for wild-type *C. elegans* embryo of *T*_cond_ = (29± 3) °C. Linear regression of *V*_in_(*T*) for individual embryos leads to the same average value of 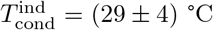 as for the ensemble (Fig. 2C, inset). With an average embryo volume *V*_embryo_ of about 30 · 10^3^*µ*m^3^ we find a relative volume of the condensed P granule phase of about 0.1 percent (see Fig. 2D; constant *V*_embryo_(*T*) see SI Fig S2). These experiments show that P granule formation in *C. elegans* embryos exhibits a well-defined condensation temperature and is consistent with thermodynamic phase separation.

### B. *In vivo* phase diagrams

Other quantities that provide important information about a thermodynamic phase separating system are the concentrations that coexist inside and outside a condensate at different temperatures. The lines connecting these concentrations define the binodals of the phase diagram.

We find that the concentrations inside and outside the P granules relax quickly to steady state values after a temperature change (SI, Sect. S2). This fast relaxation to quasi stationary concentration values suggests that P granules are at local equilibrium with their environment. More precisely, the inside and outside concentrations are at equilibrium at the P granule boundary. For temperatures between 5 to 31 °C, concentrations of PGL-1-GFP inside P granules vary between 13.2 and 5.3 *µ*M, respectively, and those outside vary between 0.37 to 0.46 *µ*M. These values imply that the partition coefficient of PGL-1 proteins decreases from 36 to 12 for increasing temperatures.

The concentrations measured inside and outside the P granules as function of temperature define the binodals of phase separation, which can be represented as a phase diagram, see Fig. 3B). The binodal on the right hand side in Fig. 3B (orange) corresponds to the inside concentration, which varies more strongly with temperature than the outside concentration described by the left hand binodal (blue). In order to understand P granule dissolution and condensation based on this phase diagram, we need to know the total average concentration *c*_total_ of PGL-1 in the cell (vertical dotted line in Fig. 3B), which was determined by immunoblotting (see SI Sect. S4). For low temperatures, the outside concentration is smaller than the total concentration. In this case the cytoplasm is supersaturated and P granules form by phase separation. At high temperatures, the local concentration outside of P granules at the P granule surface is higher than the total concentration. In this case, the cytoplasm is under-saturated and P granules dissolve. As discussed above, the outside and the total concentrations are equal at the condensation temperature of about 30 °C. These observations show that the temperature-dependent behavior of P granules in *C. elegans* can be described by a classical phase diagram. The fact that P granules dissolve upon heating suggests that this phase diagram has an upper critical solution temperature, see below.

**FIG. 3.**
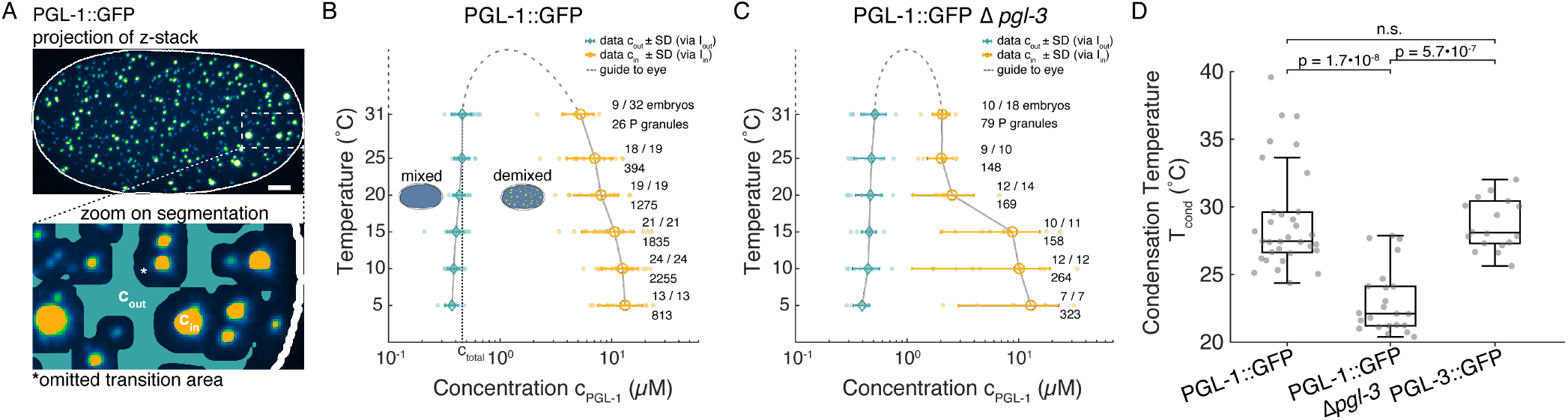
*In vivo* phase diagrams and P granule condensation temperatures. A. Illustration of the mask to determine the concentrations inside and outside the P granule condensates. To avoid the impact of the blurred condensate interface (due to point spread function and pinhole cross-talk) on the calculation of the outside concentrations, we omit a domain around the interface. We determine the concentrations inside and outside as averages over the respective domains inside and outside of the omitted regions (scale bar: 5 *µm*). B. Phase diagram for GFP-labeled PGL-1 with equilibrium concentration outside *c*_out_ (blue) and inside *c*_in_ (yellow) separating demixed and mixed states. Concentrations are calculated form fluorescence intensities *I* via a temperature corrected scaling factor between intensities and concentration (see SI Fig. S4). Number of embryos with P granules out of the total number of recorded embryos as well as the total number of P granules at each temperature are shown. Dashed line is a guide to the eye. C. Similar phase diagram as as in B but for a wormline where PGL-3 was genetically deleted which is refereed to as Δ*pgl-3*. D. Condensation temperature *T*_cond_ for different worm strains: the dissolution temperature of the wildtype with the GFP-label on the PGL-1 or PGL-3 protein is very similar. In contrast, in the case of the Δ*pgl-3* line, P granules already dissolve at approximately five degrees lower temperature.

### C. PGL-3 deletion

P granules are multi-component systems consisting of many different proteins and RNA. To shed light on the impact of the multi-component nature of P granules on their temperature response, we studied the phase diagram of P granules in a deletion line where PGL-3, one of the main protein components, was removed (see materials and methods and SI Sect. S3). We verified that the total concentration of PGL-1 stays approximately constant despite a full deletion of PGL-3 (see SI Fig. S3). The most prominent difference in temperature response of the Δ*pgl-3* line compared to the wildtype is the decrease in dissolution temperature of about six degrees to 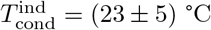 Thus, a lack of PGL-3 leads to a reduction of the temperature range of phase separation as compared to wildtype, see Fig. 3D. The phase diagram of the Δ*pgl-3* line as a function of PGL-1 concentration is shown in Fig. 3C. In the phase diagrams, we find a pronounced jump-like reduction of the inside equilibrium concentration between 15 and 20 °C. The concentrations inside decrease from 12.9 to 2.1 *µ*M while outside, concentration increases only from 0.40 to 0.52 *µ*M in the range of 5 to 31 °C. These strong changes of the equilibrium concentration inside P granules highlight the significance of PGL-3 proteins for P granule phase separation. Comparing Fig. 3B and C reveals that the presence of PGL-3 stabilizes the PGL-1 concentration inside P granules as a function of temperature in wildtype embryos. These observations are consistent with the idea that P granules form via a multi-component phase separation at local thermodynamic equilibrium. We return to this point below.

### D. Thermodynamics of *in vivo* phase separation

We propose a description for P granule phase separation based on local thermodynamic equilibrium. Quantifying intracellular phase separation using a single labelled protein component exhibits the general feature of binary phase separation resulting in a dense and a dilute phase. This phase separation can be captured by an effective free energy density *f*, which depends on the effective volume fraction of the labeled component *ϕ* and on temperature *T*. We thus introduce a binary Flory-Huggins model [15, 16] with the effective free energy per volume

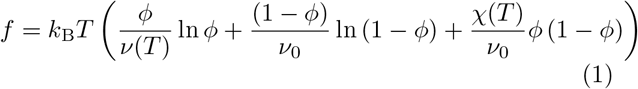

where the effective molecular volume *ν*(*T*) and the interaction strength *χ*(*T*) can depend on temperature *T*. Here, *ν*_0_ is a characteristic molecular volume of cytoplasmic molecules and the effective volume fraction is related to concentration *c* = *ϕ/ν*. The free energy density as a function of concentration given in (1) depends on temperature, as illustrated in Fig. 4A. In particular, the coexisting, equilibrium concentrations inside and outside move closer to each other when the temperature is increased and coincide at the critical point. For concentrations and temperature values outside the region enclosed by the binodal line, condensates dissolve while inside condensates are thermodynamically stable (Fig. 4A).

**FIG. 4.**
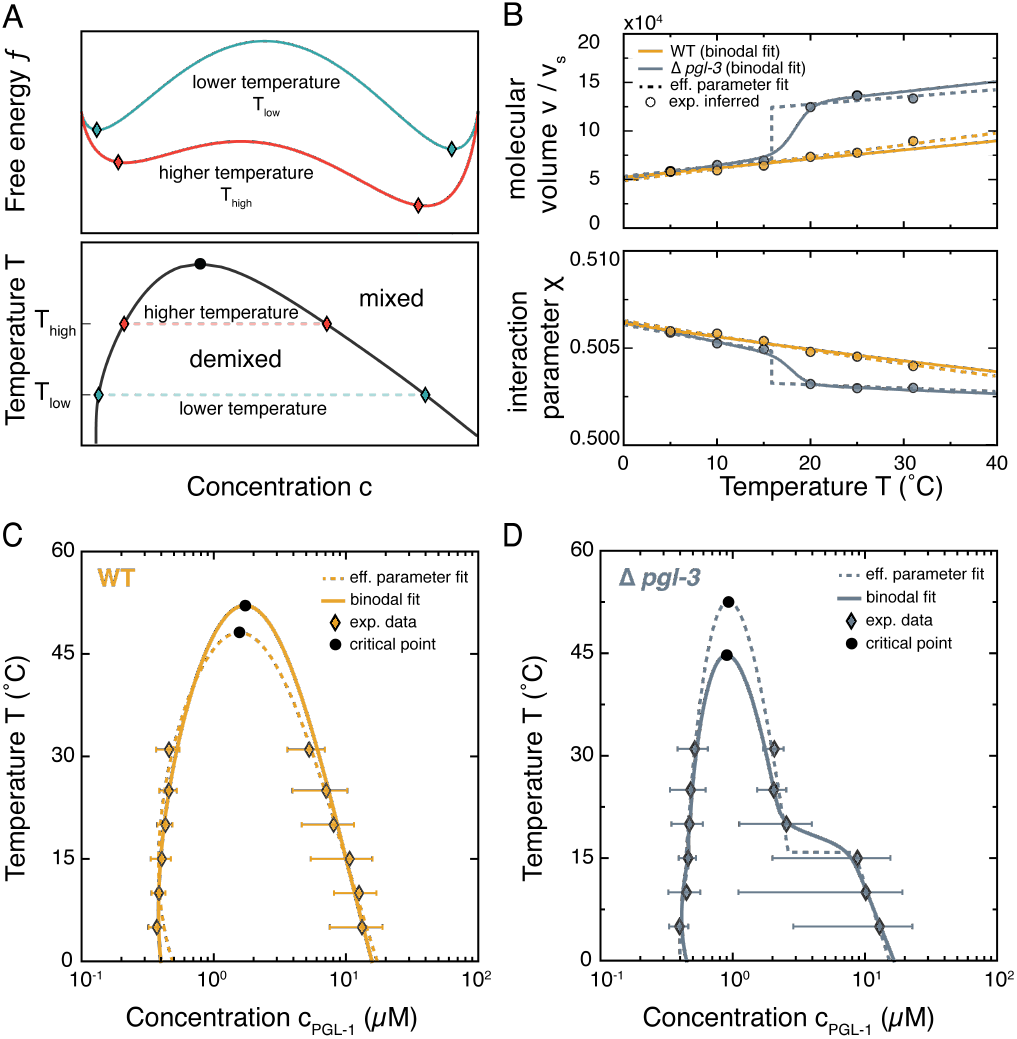
Model results for temperature-dependent phase separation. A. Free energy density as a function of concentration for two different temperatures indicating that the equilibrium concentrations inside and outside approach each other for increasing temperature (upper). This trend is also reflected in the phase diagram in which the binodal line separates the mixed and demixed phase (lower). B. From the experimental *in vivo* phase diagrams (Fig. 3C, D), we obtain via fitting the temperature-dependent interaction parameter *χ*(*T*) (orange dots) and the molecular volume *ν*(*T*) (blue dots) for each coexisting concentration pair of PGL-1 concentrations. We parameterize each function by linear functions jumping discontinuously at a temperature around 16 °C. C. and D. show the experimental phase diagram points together with the parameterized discontinuous function from C. (dashed line) and the global fit (solid line) accounting for a smoothed out discontinuity (see SI, section S1 B for details). Black circles represents the critical points.

The free energy (1) can be used to relate the experimentally observed behaviors of phase separation with the physical parameters of the Flory-Huggins model. In particular, for a pair (i.e. inside and outside a P granule) of coexisting PGL-1 concentrations measured at temperature *T* (see Fig. 3B, C), we can determine the effective interaction parameter *χ*(*T*) and the effective molecular volume *ν*(*T*) of PGL-1 using a Maxwell construction (see gray lines Fig. 4A, SI Sect. S1). The resulting temperature-dependencies of *χ*(*T*) and *ν*(*T*) are shown in Fig. 4B as circles. A key finding is that for wild-type both quantities exhibit an approximately linear dependence on temperature. Interestingly, in the Δ*pgl-3* line, the effective interaction parameter and the molecular volume exhibit jump-like changes within the interval from about 15 to 20 °C. The jump-like increase of the effective molecular volume *ν*(*T*) suggests that the PGL-1 protein changes its conformation from a collapsed to a more open, coil-like state. This transition may provide space for solvent molecules to enter the P granules leading to a reduced PGL-1 concentration (Fig. 3C).

The experimentally determined phase diagrams shown in Fig. 3B, C are limited to temperatures below the respective condensation temperatures because the average concentrations in the wildtype and Δ*pgl-3* embryo are fixed. In order to estimate the shape of the binodal line in regions where data is not available, we choose simple parametrizations of the temperature dependent interaction parameters *χ*(*T*) and the effective molecular volume *ν*(*T*) (see Materials and Methods and SI, Sect. S1 B for details). For wildtype we use linear relationships (dashed and solid orange lines in Fig. 4B). The binodal lines that correspond to these linear relationships are shown in Fig. 4C. The dashed lines are obtained from fits of linear functions to the data of *χ*(*T*) and *ν*(*T*) (effective parameter fit), while the solid lines are obtained by a fit of the calculated binodal to the concentration values in the phase diagram (binodal fit). For the Δ*pgl-3* line, we use an effective parameter fit based on a piece-wise linear dependence of the parameters, which exhibits a jump at about 16 °C (dashed lines in Fig. 4B, D). For the binodal fit we use a parameterization that connects the linear behaviors of *χ*(*T*) and *ν*(*T*) by a smooth function (see solid lines in Fig. 4B, D and SI Sect. S1 B for details). The binodal lines obtained by both the effective parameter fit and the binodal fit both follow closely the experimental data. Strikingly, the binary Flory-Huggins model can capture the experimentally determined phase separation behavior for both wildtype and Δ*pgl-3* embryos and suggests the existence of critical points at temperatures and concentrations that are outside the temperature range at which *C. elegans* can live and propagate (Fig. 4C, D). The existence of a critical point at high temperatures corresponds to an upper critical solution temperature. This suggests that the observed dissolution of P granules upon temperature increase is governed by an increase in entropy as often observed in polymer solutions [17].

The experimental data shows that the condensation temperature in the Δ*pgl-3* line is lower than in wildtype (Fig. 3D). This lower condensation temperature means that PGL-3 enhances the effective attractive interactions among PGL-1 proteins. This effect of PGL-3 is also supported by the observation that PGL-1 and PGL-3 can directly interact *in vitro* and *in vivo* [18]. Our Flory-Huggins model can be used to estimate the dependence of the condensation temperature on PGL-3 by interpolating the temperature-dependent parameters *χ*(*T*) and *ν*(*T*) between wildtype and Δ*pgl-3* line (Fig. 4C; for details see SI, Sect. S1 C). Using the wildtype PGL-3 concentration of 0.96 *µ*M, we obtain the condensation temperature *T*_cond_ as a function of PGL-3 concentration as shown in Fig. 5A. To experimentally test these predictions, we used RNA interference (RNAi) to reduce the PGL-3 concentration by approximately 60% to 0.43 *µ*M (see Material and Methods for more details). In this case, we observe a condensation temperature of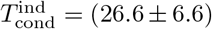, consistent with our theoretical predictions.

**FIG. 5.**
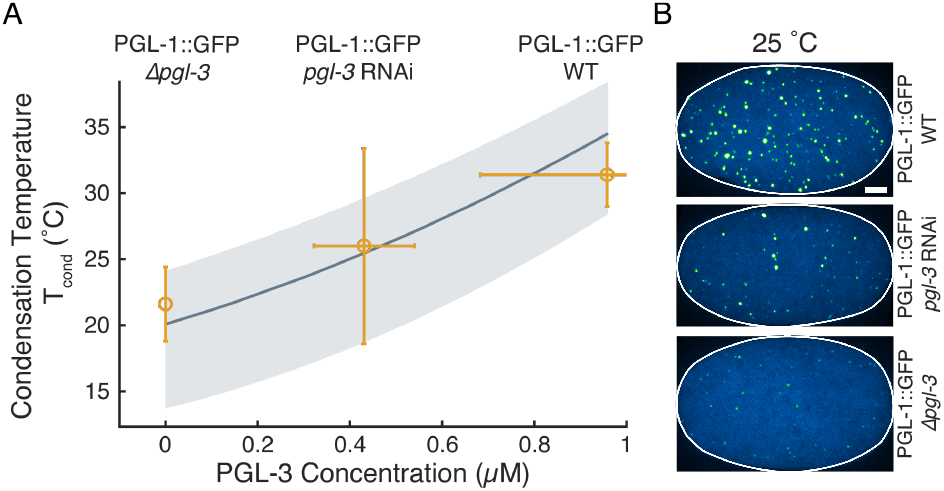
Condensation temperature *T*_cond_ as a function of PGL-3 concentration. A. The theoretical prediction is based on a linear interpolation between the fits of the experimental phase diagram data (Fig. 4C,D). While for the solid line we considered the average concentration of PGL-1, the shaded areas correspond to the lower and upper values of the standard deviation (from *in vivo* fluorescent measurements of concentrations). Experimental data are derived from linear fits to the ensemble of P granules volumes (see Fig. 2C) for WT, *pgl-3* RNAi, and Δ*pgl-3* deletion. B. Maximum projections for representative embryos at 25°C for the data presented in A. All images use the same histogram adjustments and scale bar depicts 5 *µm*.

### E. Limitations of equilibrium descriptions inside a living cell

Taken together, these results suggest that the key features of condensation and dissolution of P granules as a function of temperature are well captured by a model based on thermodynamic equilibrium. However, given that the cell is fundamentally an out-of-equilibrium system, the limitations of thermodynamic concepts need to be considered carefully. One issue is whether temperature inside the embryo is well defined since our experimental approach uses temperature shifts to probe the thermodynamic response of P granules. In this context, we can ask how fast the embryo responds to outside temperature changes. The corresponding relaxation time can be estimated based on specific heat and thermal conductivity. We find that the temperature within a size of 10 *µm* relaxes at times shorter than 100 *µs*, which corresponds to the scale of an embryo (see materials and methods for details). A corollary to this is to ask whether, at constant external temperature, heat generated inside the embryo can cause a temperature gradient towards the outside. Such heat could come from active cellular processes such as cortical and cytoplasmic flows, directed molecular transport, active force generation, and also from chemical fluxes through metabolic pathways. The heat generated by all active processes in an eukaryotic cell is typically in the order of *Q* = 10^3^ *W/m*^3^ [19–21]. For a cell of 10*µm* size, this leads to a temperature difference between cell center and periphery of only about 7 · 10^*−*7^ °C, suggesting that temperature is essentially spatially homogeneous inside the cell.

Although the temperature is essentially homogeneous in space on cellular length scales, the temperature could locally fluctuate due to active molecular processes. For example, a single ATP hydrolysis event can convert chemical energy to heat and can cause a localized and transient increase in temperature. Such perturbations relax in a diffusive manner and spread over a distance of about 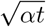 while the temperature in the corresponding volume 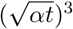 decreases with time. Here, *α* denotes the thermal diffusivity which is 1.4 · 10^*−*7^ *m*^2^*/s* in water [22]. A chemical event that releases *h*_ATP_ = 10 *k*_*B*_*T* of heat induces a temperature increase of about 10 °C after about 100 *ps* in a volume element of 1 *nm*^3^. After about 100 *ps*, this temperature perturbation has relaxed to about 10^*−*2^ °C as the released heat spreads in a volume of 10^3^ *nm*^3^. Considering even longer times we find that after 10 *ns*, the temperature perturbation has been homogenized within a volume element of diameter *d* = 100 *nm* with a remaining temperature increase of only about 10^*−*5^ °C. Furthermore, the rate at which such events are expected to occur within this volume element is given by *Qd*^3^*/h*_ATP_ ≃ 25 *s*^*−*1^ and is slow compared to the temperature relaxation time of 100 *µs*. Thus, at longer time scales, we can consider volume elements of size 100 *nm* to be at local thermodynamic equilibrium and we can neglect temperature fluctuations arising from individual chemical events.

## III. DISCUSSION

### A. Local thermodynamics and the non-equilibrium nature of the cell cytoplasm

Work over the last decade has accumulated evidence that the formation of intra-cellular condensates can be described by the physics of phase separation. However, it has been unclear whether thermodynamic concepts are appropriate to explain condensate formation despite the presence of numerous active processes in living cells. Here we have shown that P granule formation and dissolution in the *C. elegans* embryo has a temperature dependence as expected for a thermodynamic process and can be represented by a classical phase diagram. Our analysis demonstrates that the P granule dynamics are governed by local thermodynamic equilibrium in mesoscopic volume elements of about 100*nm* size beyond time-scales of 1*µs*. Estimates based on general arguments show that temperature heterogenities due to individual active molecular events dissipate quickly without perturbing the slower dynamics of phase separation. Consistently, a simple phase separation model based on thermodynamics can well describe the phase diagrams determined *in vivo*.

Condensates in cells are typically composed of many different molecular components which interact with each other. Here, we observe the condensation of one component, PGL-1. However, the resulting phase diagram should depend on the concentrations of other main components. Indeed, when we deleted PGL-3 proteins in the *C. elegans* embryo (Δ*pgl-3* line), we found a qualitative change of the phase diagram reflected in the appearance of a jump-like change of the PGL-1 concentration inside P granules with temperature. We do not know the origin of this jump like response. However, it could reflect a signature of another phase transition in the multi-component system. This transition may be associated with conformational changes of PGL-1 proteins as suggested by the jump in molecular volume that we infer in our analysis. Our results suggest that the presence of PGL-3 suppresses this conformational transition. More generally, our results suggest that the multi-component nature of condensates helps to protect the condensate’s behavior and the proteins within it against changes in physiological conditions and other stresses. For instance, *C. elegans* propagates in an appropriate range of temperatures of approximately 10 °C to 30 °C degrees and condensates must be able to operate effectively within such changes.

Temperature changes can also affect rates of chemical reactions relevant to P granule formation. For example, DEAD-box ATPases [7] were shown to promote phase separation in their ATP-bound form, whereas ATP hydrolysis induces compartment turnover and release of RNA [8]. However, such chemical rates typically vary only little. For example, enzymatic reactions change at most 2-3 fold within a temperature range of (20 − 30) degrees [23]. Similarly, in the same temperature range, rates of active processes driven by the hydrolysis of ATP or GTP change also only weakly with temperature [24, 25]. Together with the excellent agreement between the *in vivo* phase diagrams and the equilibrium phase separation model, we conclude that the temperature dependence of rates associated to active processes do not play a key role in the temperature response of P granules.

We have proposed that volume elements of size of about 100*nm* are locally at equilibrium, which is why local thermodynamics can be used to describe phase separation of P granules in the *C. elegans* embryo. How can we relate the small scales that are at equilibrium to larger scales in the cell where active dynamic processes take place? The non-equilibrium active character of the cell typically emerges on length scales larger than the size of this volume element. For example, concentration levels change in time, concentration profiles develop, and active transport processes and fluid flows stir the inside of living cells. Eventually, heat is released to the cell environment. The thermodynamic driving forces of such cellular processes are for example differences in chemical potential between different volume elements that are in different local equilibria. It is important to remember that a local equilibrium means that local volume elements relax to an equilibrium state quickly, faster than the time scales during which concentrations and thermodynamic variables change. However, at longer time scales, individual volume elements can change their properties because they are in general not at equilibrium with their environment. This is why, in living cells, non-equilibrium phenomena can be captured by classical non-equilibrium thermodynamic approaches that are built on local thermodynamics [11, 26, 27].

It is interesting to estimate the actual chemical activity in such local volume elements. Taking into account the estimated rate of heat production per volume and the energy released as heat during individual chemical events, we estimate on average about 25 active chemical events per second in a volume element of 100 *nm* size. This is much slower than the microsecond relaxation time to equilibrium and therefore does not significantly perturb the local equilibrium. However, it provides the basis for the non-equilibrium phenomena that emerge at larger scales inside the cell (see materials and methods for an estimate of the length scale relevant for the example of protein expression). As a result, spatial heterogeneities emerge because different volume elements are not at equilibrium with respect to each other. It is important to note that the simplified picture based on 100 *nm* volume elements outlined here applies to a well mixed and isotropic cytoplasm. Inside or near membranes or other cellular assemblies, the construction of volume elements will have to be refined.

### B. Possible role of thermodynamics for organism fertility

So far we have shown that the temperature-dependent response of P granules in the *C. elegans* embryo is governed by local thermodynamics. It is interesting to relate the temperature dependence of P granules to the temperature dependence of fertility. The fertility of wildtype worms has been reported to describe a bell shaped curve for temperatures between about 10 and 27 °C [14] with a maximal fertility at about 18 °C. Interestingly, fertility is lost at about 27 °C when P granules have largely dissolved, see Fig. 2D and Fig. 3 in Ref. [14]. This observation suggests a potential link between fertility at higher temperatures and P granule thermodynamics. Indeed, it has been reported that worms deficient in PGL-1 are more temperature sensitive than wildtype and further need PGL-3 for fertility at lower temperatures [18].

Interestingly Δ*pgl-3* worms have a temperature range of fertility that is slightly shifted to lower temperatures (Fig. S5). Our data show that this is correlated with a decrease in condensation temperature, see Fig. 3D and Fig. S6. Thus, in Δ*pgl-3* worms, the temperature range where P granules are abundant is shifted to lower temperatures. This shift suggests that the multi-component nature of P granules may help to modulate the temperature range of fertility during evolution. Note that when comparing the fertility of N2 worms and Δ*pgl-3* mutants in a N2 background, we observed a higher brood size of the mutants at lower temperatures of 10°C (see SI Fig. S5) for reasons we do not yet understand. However, we see a possible link between total condensate volume and condensation temperature *T*_cond_ with the fertility of the worm, suggesting further exploration of the thermodynamic properties of *in vivo* systems is needed.

## IV. MATERIALS AND METHODS

### A. *In vivo* temperature perturbations

From Fig. 2 onward, early one-cell *C. elegans* embryos before the onset of P granule segregation were used. *C. elegans* strains were kept and cultured at 16 °C on standard feeding plates. Prior to measurement, adult worms were dissected in M9 buffer containing 25 *µm* diameter polystyrene spacer beads (Polysciences, USA) under temperature controlled conditions. A drop of 8.1 *µl* of this solution containing embryos of various developmental stages was sandwiched between a 18 by 18 mm coverslip and the sapphire glass of a custom-made temperature stage (as previously described in [28] and see sketch in SI Fig. S1). The chamber was sealed using a silicone glue (twinsil-speed, Picodent, Germany) ensuring no evaporation occurred. For imaging we used a CSU-X1 (Yokogawa, Japan) spinning disc confocal system on an IX83 microscope (Olympus, Japan) connected to an iXon DU-897 back illuminated EMCCD camera (Andor, UK). All experiments were acquired with 0.7 *µm* z-spacing using an objective piezo (PiezoJena, Germany) with a 40X UPLSAPO 0.95 NA air objective (Olympus, Japan). Emission laser intensities were kept constant and measured prior to imaging to ensure comparability throughout the entire data-set. The microscope was controlled via VisiView (Visitron Systems, Germany). The z-stacks in Fig. 2 A were captured every 30 s, for other experiments a protocol was used as described in SI Fig. S1. In brief, every embryo was subjected to 4 temperatures in the range of 5 to 25 °C where 3 min were allocated after each temperature change to reach local equilibrium (Control for timing see SI Fig. S7. All embryos where subjected to 31 °C in the final step to allow for relating fluorescence values between all worm strains and experiments. This is facilitated since most P granules are already dissolved at this temperature. In worm gonad images shown in Fig. 1 B were imaged using a CSU-W1 (Yokogawa, Japan) spinning disc confocal system on an IXplore IX83 microscope (Olympus, Japan) with a 40X UPLSAPO 0.95 NA air objective (Olympus, Japan) controlled via CellSens. Worms were mounted like described previously for embryos, however, using 50 *µm* diameter polystyrene spacer beads.

### B. Image registration and *in vivo* concentration determination

Image registration and analysis was performed using a custom-written MATLAB pipeline (The MathWorks, USA). Image stacks were both analyzed in 3-dimensions and as 2D maximum and mean projections. All image stacks were first subtracted by the dark intensity count of the camera. Both 2D and 3D images are subjected to filtering prior to registration of condensates and embryo contour. For 3D analysis the Gaussian blur kernel is also adopted to represent different resolutions in planar (0.125 *µm*) and axial direction (0.7 *µm*). The shape of the embryo outline is detected and used for total intensity *I*_total_ and solute intensity calculation, the embryo volume, as well as further processing. P granule outlines are registered via a threshold using the embryo mean intensity + a multiple of its standard deviation (condensate volume detection see Fig. 2B, *c*_in_ and *c*_out_ mask see Fig. 3A). For the determination of *c*_in_ via the condensate intensities a higher threshold than for the determination of the volumes *V*_in_ was used. This suppresses the influence of very small condensates that have unrepresentative intensities for the quantification of *c*_in_. In addition, a watershed procedure was used to separate falsely connected condensates before estimating their volume. To determine *c*_out_ via the solute intensity, the mask of the *V*_in_ was dilated and then subtracted from the registered embryo outline which left the solute mask. The dilation leaves a gap to the mask used for the determination of *c*_in_ (see Fig. 3A). This reduces the influence of the tail of the Gaussian-like profile of condensate intensity-distributions and possible pinhole cross-talk of the spinning disc system on the solute intensities. The relation of intensity values to concentration was set via a temperature dependant conversion factor *α*_T_ using the assumption of a linear relationship (see SI Fig. S4). To get *α*_T_, the average total intensity *I*_total_ values obtained from registration of whole embryos and the average total concentrations *c*_total_ determined via quantitative western blots for each worm strain were related and then pooled for each temperature (see SI Fig. S3). That this linear relationship is reasonable can be seen when comparing the average *I*_total_ and the average *c*_total_ between worm strains, where we report comparable values for both approaches (see SI Fig. S3).

### C. RNAi experiments

A colony of bacteria from a ampicillin (100 *µg l*) feeding plate was cultivated over night in LB medium containing ampicillin and tetracycline at 37 °C at 220 rpm on shaker plate. After centrifugation (4000 rpm, 5min) the pallet was re-suspened to an OD of 0.5 in LB medium containing ampicillin and 0.2mM IPTG. After another 2 h at 37 °C at 220 rpm 150 *µl* were pipetted on a dryed NGM feeding plate and either used directly or stored at 4 °C in a dark environment. Induction of RNAi was carried out with L4 stage worms for 24 h at a temperature of 23 °C [29]. For *pgl-3* RNAi both worm strains expressing PGL-3::GFP and PGL-1::GFP were cultivated under the same conditions at the same time. While experiments were carried out with the strain expressing PGL-1::GFP, the PGL-3::GFP strain served as means to measure the amount of reduction in PGL-3 via the relation of total fluorescence 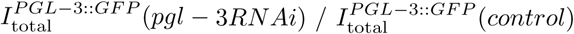 (see SI Fig. S8A). The relative total intensity 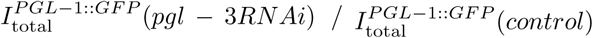 for PGL-1::GFP was not significantly changed using *pgl-3* RNAi (see SI Fig. S8B).

### D. *C. elegans* worm strains

The proteins PGL-1 and PGL-3 are regarded as regulators of the germ line and belong to the main constitutive components of P granules [30]. To experimentally analyze the phase separation behavior of P granules, their main components PGL-1 or PGL-3 were labeled (tagged) at their endogenous genomic locus with monomeric enhanced GFP (mEGFP) using the co-CRISPR method [31]. The following lines were used: N2 wildtype worms; TH586: *pgl-1* ::mEGFP pgl-1(dd54[pgl-1::mEGFP]); TH561: *pgl-3* ::mEGFP (pgl-3(dd29[pgl-3::mEGFP]); TH615: ∆*pgl-3* (dd47) pgl-1::mEGFP (pgl-1(dd54[pgl-1::mEGFP])); TH605: ∆*pgl-3* (dd47); TH721: ∆*pgl-1* (dd65) pgl-3::mEGFP (pgl-3(dd29[pgl-3::mEGFP]) (see SI, section S3 for details on the analysis).

### E. Protein concentration determination

Previously, protein concentrations were measured in N2 wildtype early embryos by quantitative label free mass spectrometry [32]. For measuring protein concentrations of PGL-1 and PGL-3 in the worm lines used in this study, PGL-1 and PGL-3 were specifically detected in immunoblots of early embryo protein extracts with monoclonal antibodies and imaged with an Odyssee SA Licor infrared imaging system. The protein concentrations were then calculated by normalizing to the wild-type values. Protein concentrations were measured from 5 independent experiments (see SI, section S4 for details on the analysis).

### F. Fit of a binary mixture model to the experimental data

Coexisting phases in incompressible binary mixtures satisfy the following conditions [33],

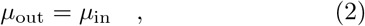

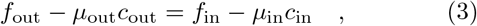

where *µ* denotes the chemical potential defined by *µ* = *∂f/∂c* and the subindices in and out indicate the two distinct phases. The conditions given in Eqs. (2) and (3) represent the balance of exchange chemical potentials and osmotic pressures between the phases, respectively. We numerically solve these equations and find for each coexisting concentration pair the interaction parameters *χ* and *ν* leading to the best match with the experimentally determined values. These set of values for *χ* and *ν* are shown as dots in Fig. 4 B. From these results, we obtain the temperature dependence of *χ* and *ν* which in the case of the ∆*pgl-3* line shows a transition occurring between 15 and 20 °C. We then fit curves for each corresponding set of points (dotted lines in Fig. 4 B) and calculate the corresponding phase diagrams (dotted lines in Fig. 4 C,D) which agree very well with the experimental phase diagram. For details on the specific form of the functions we refer the reader to SI S1 B. In addition, to further assess the dependence of these parameters as a function of temperature we also study the temperature dependence of the parameters by minimizing a global error function defined by

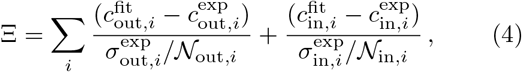

where each index *i* corresponds to a different temperature, and 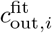 and 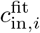 are the values obtained from solving the equilibrium conditions (S2) and (S3). Moreover, 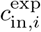 denote the experimentally measured values, 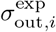 and 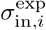 are the standard deviations of the measurements and 𝒩_out,*i*_ and 𝒩_in,*i*_ correspond to the number of data points for each temperature. The solid lines in Fig. 4B show the curves that minimize the global error (4) and the corresponding phase diagrams are shown as solid lines in Fig. 4C,D. We find that both approaches are in very good agreement with the experimental phase diagrams. In order to estimate the condensation temperature of the P granules as a function of PGL-3 concentration, we construct phase diagrams corresponding to different concentrations of PGL-3, where we choose the dependence of the parameters defining *χ* and *ν* as a linear interpolation between the values of the interaction parameter and the effective molecular volume of the phase separating component corresponding to the wildtype and the ∆*pgl-3* line. For this interpolation, we consider the parameter values corresponding to the wildtype line obtained by the global fit (solid yellow line in Fig. 4 B) and for the ∆*pgl-3* line we only consider the points below the transition temperature, i.e. the parameters obtained from the fit to the coexisting phases corresponding to temperatures 5, 10 and 15 °C. Using these considerations we estimate the condensation temperatures as a function of PGL-3 concentration for a fixed concentration of PGL-1 (value from immunoblotting), we repeat the same process for concentrations that are one standard deviation above and below this estimate. The predictions shown in Fig. 5 as the grey shaded area are in very good agreement with the experimental data.

### G. Estimation of temperature relaxation times, temperature gradients and temperature fluctuations

In this work we characterize phase separation as a function of temperature in a living cell. A living cell is maintained at all times far from equilibrium, for example due to the generation of heat by metabolic processes and the production and degradation of molecular components. This non-equilibrium nature raises the question whether temperature is a reliable thermodynamic variable that can be controlled experimentally. This question involves how fast temperature changes can be applied and what are stationary temperature difference inside cells due to metabolic processes.

#### Temperature relaxation times in the embryo

In a typical *in vitro* assay, an experimenter can wait until the system has relaxed toward thermodynamic equilibrium before taking a measurement. In measurements on cells, however, no thermodynamic equilibrium is reached and the time window to take the measurement is limited because the cell cycle progresses. In order to characterize the temperature dependence of cellular processes, the minimal time between measurements at different temperatures should be larger than the temperature relaxation time. The time dependence of the temperature profile is described by the heat equation

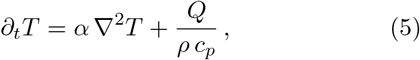

where *α* denotes the thermal diffusivity, *ρ* is the mass density, *c*_*p*_ specific heat capacity at constant pressure, and *Q* denotes the volumetric heat source due to metabolic processes. A temperature perturbation of size 𝓁 relaxes on a time-scale *τ*_*T*_ (𝓁) = 𝓁^2^*/*(6*α*). For the embryo, we estimate the thermal diffusivity in the embryo similar to water with *α* = 0.14mm^2^*/s* [22]. *Considering an embryo size of* 𝓁 ≃ 10*µm*, the time to relax temperature heterogeneities is *τ*_*T*_ *≃* 100*µs*. In our experiments (Fig. S1), heat is exchanged via sapphire with a thermal diffusivity of about 100*α*. Therefore, contributions to the temperature relaxation of sapphire can be neglected. In summary, for processes that occur on time-scales exceeding *τ*_*T*_, such as the P granule growth occurring on second time scales, temperature has relaxed and takes stationary values inside the embryo.

#### Effects of heat generation due to metabolic processes

Exothermic metabolic processes generate heat inside the embryo which is conducted to the embryo boundaries. The heat production is of the order of 1 *W/kg* for humans [19] as well as for flat worms [21]; for a rather complete overview of metabolic rates corresponding to a large number of species, see Ref. [20]. Here, we consider the heat production of the flat worm as a good estimate for the *C. elegans* embryo. Using a mass density of 10^3^*kg/m*^3^ the corresponding volumetric heat source is *Q* ≃ 10^3^*W/m*^3^. In steady state, this heat source creates a temperature profile. For an embryo of width *L* = 20*µm* that is squeezed between two heat conducting surfaces located at *x* = ±*L/*2 and maintained at ambient temperature *T*_ext_, the temperature profile reads

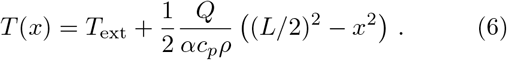

This equation implies that the stationary temperature difference relative to the ambient temperature are only *T* (*x* = 0) – *T*_ext_ ≃ 7 · 10^*−*7^ °C using the specific heat *c*_*p*_ = 4 · 10^3^*J/*(*kg K*) of water. These weak temperature differences allow to consider temperature as a roughly constant thermodynamic field in the embryo.

Finally, we estimate the impact of temperature fluctuations due to exothermic metabolic events. We consider that a single event dissipates an energy *h*_ATP_ and occurs at a position 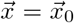. For this case, the volumetric heat source reads

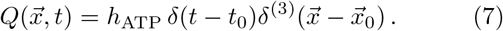

The corresponding temperature profile for the propagation of this single metabolic event in time and space (ignoring boundary effects) reads

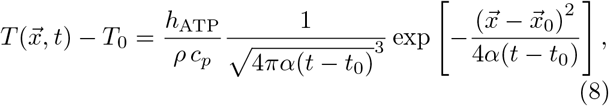

for *t* > *t*_0_, where *T*_0_ is the embryo temperature prior to the metabolic event.

To estimate the role of temperature fluctuations in the embryo, we consider the time for a single metabolic event to diffusive through a volume of size *d, τ*_*T*_ = *d*^2^*/*(6*α*), and estimate the corresponding temperature change. In particular, according to (8), the temperature change ∆*T* (*d*) in the volume element of size *d* due to a single metabolic event is given by ∆*T* (*d*) = *h*_ATP_*/ ρ* (*c*_*p*_*d*^3^). Using for the mass density *ρ* and for specific heat *c*_*p*_ the value of water, and *h*_ATP_ = 10*k*_*b*_*T*, we find that ∆*T* (*d*) = 10 °C · (1 *nm/d*)^3^. The corresponding time-scale is *τ* (*d*) *≃* 1 *ps* · (*d/*1 *nm*)^2^. These estimates suggest that for volume element above sizes of *d* = 100 *nm*, corresponding to time-scales above 10^*−*8^ *s*, temperature changes due to single metabolic events are less than 10^*−*5^ °C and thus negligible.

#### Effects of sources and sinks of molecular components

In general, molecular components such as proteins inside cells are not conserved; they are continuously produced and degraded with time. Cells can increase or decrease the concentration of specific proteins by regulating the protein expression machinery. The corresponding reaction cycles in general include multiple feedback loops which give rise to a characteristic rate of change *k* of the protein concentration. This rate is typically small and in the order of *k* ∼ 1(h)^*−*1^ or even slower [34]. To estimate the impact of sources and sink of molecular components, the relevant quantity to be estimated is the reaction-diffusion length scale 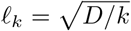, where *D* denotes the diffusion coefficient of the protein. Above this length scale, protein expression will affect the kinetics of phase separation [11]. Below this length scales, the behavior of the system is expected to be governed by local thermodynamics, at least with respect to the protein expression kinetics. Specifically, using typical protein expression rates given above and an approximate PGL-1 diffusion constant of *D* ≃ 1*µ*m^2^*/s* [35], *gives 𝓁*_*k*_ ≃ 60*µm*. Interestingly, this length scale is approximately equal to the size of the embryo. Thus, protein expression can affect phase separation on the embryo scale, however has not much effects on the local thermodynamics governing the formation and dissolution of P granules.

## ACKNOLEDGEMENTS

We would like to thank Shambaditya Saha, Louise Jawerth, Marcus Jahnel, Titus Franzmann, Juan Iglesias, Patrick McCall, Tyler Harmon, and the whole Hyman-lab for fruitful discussions about phase separation during the last years, and Melissa Rinaldin and Jonathan Rodenfels for very helpful feedback on the manuscript. Falk Elsner and Hartmut Wolf for support in optimizing the temperature stage. We also thank Susanne Ernst and Anne Schwager for technical assistance with worm work and immunoblotting and Torsten Bücher for technical assistance with brood size measurements. We thank Olympus for providing the spinning-disc system based on an IXplore IX83 microscope that was used for whole worm imaging. A.W.F. was supported by the ELBE postdoctoral fellows program and the MaxSynBio consortium, jointly funded by the Federal Ministry of Education and Research of Germany and the Max Planck Society. O.A.A. acknowledges funding from the Armando and Maria Jinich postdoctoral fellowship for Mexican citizens. C.A.W. and A.A.H. acknowledge the SPP 2191 “Molecular Mechanisms of Functional Phase Separation” of the German Science Foundation for supporting the exchange among scientists working of intra-cellular phase separation.

## AUTHOR CONTRIBUTIONS

A.W.F. performed all *in vivo* measurements shown in this work. Experiments are based on the findings of A.F.D.-D.’s thesis work on temperature response of P granules. A.F.D.-D. and A.A.H. initiated the project. A.W.F., O. A.-A. and C.A.W. developed the data analysis pipeline. O.A.-A., A.W.F., F.J. and C.A.W. worked on the theory and the fitting to the experimental data. C.H. performed immunoblotting and prepared the CRISPR worm lines. M.M. and M.K. contributed to the expertise of cellular temperature perturbations and the configuration of a spinning-disc setup. M.L. and C.H. performed and analysed the brood size and the fertility measurements. A.W.F., A.F.D.-D., O.A.-A., C.H., M.L., A.A.H., F.J. and C.A.W. conceived the project and wrote the paper.

## SUPPLEMENTAL INFORMATION

### S1. THERMODYNAMIC MODEL

#### A. Definition of effective binary model and equilibrium conditions

As discussed in the main text, we model the cytoplasm as an effective binary mixture composed of a phase-separating component which can form P granules and a solvent component that accounts for the rest of the cytoplasmic liquid. The phase-separating component encompasses all the different proteins and biomolecules that localize into P granules, in particular, we use the concentration of PGL-1 as a probe of this component and include the effects of the rest of the constituents using an effective molecular volume *ν*(*T*) and an effective interaction parameter *χ*(*T*), which in general depend on the temperature and composition of the mixture. In order to quantitatively study the binary mixture we define a free energy density *f* of the form:

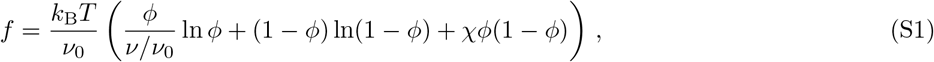

where *k*_B_ is the Boltzmann constant, *T* is the absolute temperature, *φ* is the volume fraction of the phase separating component and the effective molecular volumes of the phase separating component and the cytoplasmic liquid are *ν* and *ν*_0_, respectively. Incompressibility can be satisfied when considering that changes in protein molecular volume with temperature dominantly occur via uptake or release of cytoplasmic molecules. The first two terms on the right hand side of (S1) represent the mixing entropy of the system, while the last term describes the tendency of the system to form two distinct phases. The explicit dependence of (S1) in the concentration of PGL-1, denoted by *c*, follows from the relation *φ* = *ν c*.

The homogeneous exchange chemical potential of the phase-separating component, *µ* = *ν∂f/∂φ*, is used to express the coexistence conditions,

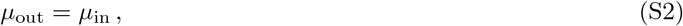

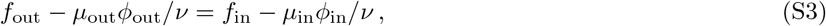

stating that the exchange chemical potential and the osmotic pressures of the two coexisting phases (inside (‘in’) and outside (‘out’)) balance at thermodynamic equilibrium. The conditions above relate each temperature to an unique pair of consecrations corresponding to the coexisting phases inside and outside P granules.

#### B. Fitting of thermodynamic model to the experiments

We obtain interaction parameter *χ*(*T*) and the effective molecular volume of the phase-separating component *ν*(*T*) by fitting the experimentally determined concentrations of the coexisting phases shown in the phase diagrams Fig. 3 B,D. In particular, we fix the temperature and numerically solve (S2) and (S3). By varying the interaction parameter and the molecular volume we search for best match of the numerically calculated concentrations *c*_out_ and *c*_in_ with the corresponding experimental values. The corresponding results for *χ*(*T*) and *ν*(*T*) are depicted in Fig. 4 B. This procedure amounts to representing the information contained in the experimental phase diagrams in terms of system-specific and temperature-dependent values for the interaction parameter *χ*(*T*) and the molecular volume *ν*(*T*). Interestingly, the concentration jump observed in the experimental phase diagram for the ∆pgl-3 line (Fig. 3 C) is also observed in both parameters as a function of temperature as shown in Fig. 4 B.

This new representation of the experimental data allows us to suggest a parametrization of the temperature dependence of both the interaction parameter and the molecular volume in terms of piece-wise linear functions which account for the observed jump-like behavior:

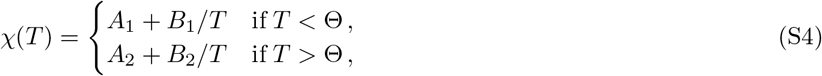

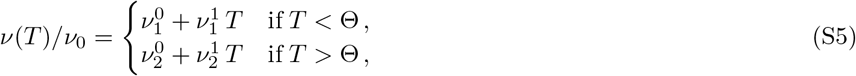

where 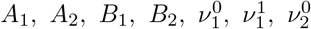 and 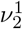 are fitting parameters. Moreover, Θ is the transition temperature at which *ν*(*T*) and *χ*(*T*) sharply change and is located between 15 °C and 20 °C. We choose *θ* =16°C for the all fits; see Fig. 4 B. We use (S4) and (S5) to determine the fitting parameters characterizing *χ* and *ν/ν*_0_ by fitting the corresponding experimental phase diagrams; these fits are shown as dashed lines in Fig. 4 B. Using the parameters obtained from the fits, we compute the phase diagram of the effective binary mixture, which are shown as dotted lines in Fig. 4 C,D, and find very good agreement with the experimental results.

The parametrization of the molecular interactions and effective volumes in (S4) and (S5) has the disadvantage of creating a potentially artificial, step-like behaviors. The transition might in principle be smooth. Hence in order to obtain a continuous smooth fitting function for the interaction parameter and the molecular volume ratios we draw inspiration from (S4) and (S5) (in terms of the transition temperature Θ and the linear slopes) and propose the following smooth functions:

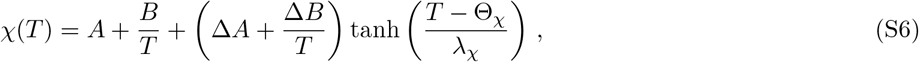

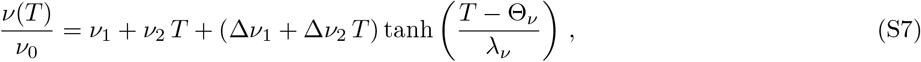

where *A, B*, Δ*A*, Δ*B, θ*_*χ*_, *λ*_*χ*_, *ν*_1_, *ν*_2_, Δ*ν*_1_, Δ*ν*_2_, *θ*_*ν*_, *λ*_*ν*_ are all fitting parameters. In order to obtain the global fit we minimize the following error function

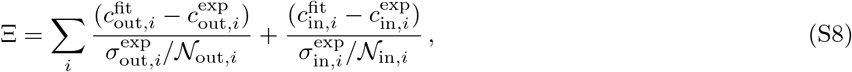

where each index i corresponds to a different temperature, and 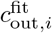 and 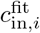 are the values obtained from solving the equilibrium conditions (S2) and (S3). Moreover, 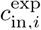 denote the experimentally measured values, 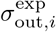 and 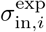 are the standard deviations of the measurements and _out,*i*_ and _in,*i*_ correspond to the number of data points for each temperature. Finally, for the case of the Δpgl-3 line, in order to avoid a large bend of the binodal, we introduce a pair of coexisting phases between the experimentally measured values of the concentrations at 15 °C and 20 °C. We choose these intermediate concentrations to be average of the concentration values of the dilute and condensed phases at 15 °C and 20 °C. We use a weight in the fitting procedure which is the average of the weights associated to the dilute and condesed phases at 15 °C and 20 °C.

#### C. Modeling the effects of PGL-3 concentration on phase separation

We assess the effect of changes in PGL-3 concentration in the P granule condensation temperature by assuming a linear extrapolation from the set of parameters obtained from the global fit to the phase diagram corresponding to the WT to those corresponding to the Δ*pgl-3* line in the piece-wise approximation in the region below the transition temperature. In order to obtain the condensation temperatures, we fix the PGL-1 concentration and estimate the parameter values as a function of the PGL-3 concentration *c*_*pgl-3*_ using the following functions

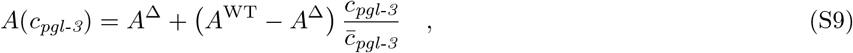

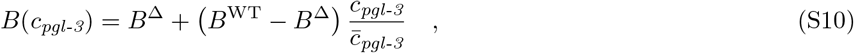

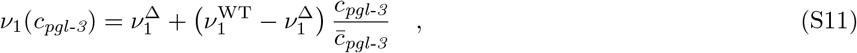

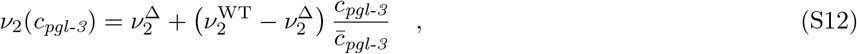

where the superindices WT and Δ indicate the parameters corresponding to the different worm-lines and 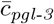 is the PGL-3 concentration estimated from immunoblotting experiments. Using the simple functional forms (S4) and (S5) and the linear extrapolation functions (S9)-(S12), we construct the corresponding phase diagrams and determine the condensation temperature for three different PGL-1 concentrations corresponding to the the mean value estimated from immunoblotting and the concentrations corresponding to deviations of one standard deviation above and below the mean value, where the standard deviation is taken from the fluorescence measurements.

### S2. EXPERIMENTAL WAITING TIME TO APPROXIMATELY REACH LOCAL THERMODYNAMIC EQUILIBRIUM

The time window for experiments in an early one-cell embryo is only approximately 15 min and we tried to fit as many temperature steps into one experiment as possible. We carried out a control experiment to estimate the time until P granules reach a local thermodynamic equilibrium. We capture a series of z-stack according to the protocol of the actual measurement (0.7 *µm* z resolution) in intervals of 30s and apply a temperature step. We choose a step of 10 °C between 15 and 25 °C. From the data we collected, we conclude that a waiting time of 3 min is enough to ensure a settled response of both P granule volume and numbers (see S7). This lead us to the design of the protocol we used throughout the manuscript (see S1).

### S3. WORM LINES AND SAMPLE PREPARATION

#### A. Generation of worm lines by CRISPR

For gene tagging by CRISPR, N2 wildtype young adults were injected with a protein/RNA mix (4.3 *µM* Cas9, PEP facility MPI-CBG; 33 *µM* tracRNA, 24 *µM* cRNA, Dharmacon; 1*µg/µl* PCR purified GFP tagging cassette; cRNA dpy-10, 9 *µM* ; recombination oligo dpy-10, 1.25 *µM*, IDT) which was assembled for 15min at 37°C, and filtered through spin columns (Durapore PVDF, 0.22 *µm*, Millipore) and grown at 20°C for 4 days. F1 animals were screened for roller (dpy-10) phenotypes and singled to new plates[31]. Homozygous F2 animals with GFP expression were identified by fluorescence screening. Worm lines were tested by fluorescence microscopy and immunoblotting for correct expression. TH586 animals were injected with protein/RNA mix (as above, but with two cRNA targeting pgl-3, each 24 *µM*, Dharmacon and a recombination oligo targeting the full ORF of pgl-3, 10 *µM*, IDT) targeting pgl-3 and grown at 20°C for 4 days. F1 animals were screened for roller (dpy-10) phenotype. Δ*pgl-3* deletion animals were identified by PCR and homozygous Δ*pgl-3* worms recovered after backcrossing to N2 males and PCR screening. Δ*pgl-3* genotype of homozygous progeny was confirmed by DNA sequencing and immunoblotting.

##### Worms lines used

- N2 wt [**?**]: received from Caenorhabditis genetics Center, CGC.
- TH586: PGL-1::mEGFP pgl-1(dd54[pgl-1::mEGFP]). cRNA 26 was targeting 5’GGGGGTCGTGGTG-GACGCGG3’; PCR tagging was done with the oligo CH578 5’GAGATCGTGGACGTGGTGGT-TACGGGGGTCGTGGTGGTCGTGGCGGTTTCAGTAAAGGAGAAGAACTTTTCACTGGAGTTG3’ creating a C terminal mEGFP fusion without any additional amino acids.
- TH561: PGL-3::mEGFP (pgl-3(dd29[pgl-3::mEGFP]). cRNA 3 was targeting 5’GAAGGCTTAGGAACCTC-CAC3’. PCR tagging was done with primer 5’GACGTGCCGGATTCTTTGGTGGATCCCGTGGAGGTTCCGAAGTGCATACCAATCAGGACCCGC3’ creating a C terminal mEGFP fusion with 2xTY linker (EVHT-NQDPLDEVHTNQDPLDTS) in front of the mEGFP.
- TH605: Δ*pgl-3* (dd47). Full deletion of pgl-3 by CRISPR with recombination oligo CH678 5’CAAACGACAAATTGTGGAGGTCGATGGGATTAAGTCCTACTTTTTCCCCCAGGGAG-GACGTGCCGGATTCTTTGGTGGATCCCGTGG3’
- TH615: Δ*pgl-3* (dd47) PGL-1::mEGFP (pgl-1(dd54[pgl-1::mEGFP])) (Δ*pgl-3* was introduced with oligo CH678 5’CAAACGACAAATTGTGGAGGTCGATGGGATTAAGTCCTACTTTTTCCCCCAG GGAGGACGTGC-CGGATTCTTTGGTGGATCCCGTGG3’)

#### B. Worm growth and embryo extract preparation

Worm lines were grown at 16°C on NGM OP50 plates with 3.6% agar and starved to dauer larvae. Dauer larvae were washed off and released onto fresh plates and grown synchronously until young adults started egg laying. Worms were washed off and embryos harvested by vigorous shaking in bleaching solution (20% sodium hypochlorite, 200mM NaOH) and several washes in M9. Embryo solutions were counted, pelleted and extracts prepared by addition of HU buffer (8M Urea, 5% SDS, 200mM Tris pH 6.8, 1mM EDTA, 1.5% DTT) and shaking at 65°C for 15min. Extracts were intermittently sonicated for 10s with a microtip after 5min incubation at 65°C to fully lyse embryos. For each worm line (N2 WT, TH586, TH615, TH561, TH721) five independent extracts were prepared from independent worm cultures and frozen at -20°C.

### S4. IMMUNOBLOTTING AND QUANTIFICATION OF PGL PROTEINS

Embryo extracts were separated on 4-12 % NuPAGE Bis-Tris gels (Thermofisher) and blotted for 20 min onto Nitrocellulose (iBlot, Thermofisher). Total protein was visualized by revert protein stain [**?**] and imaged at 700 nm in an Odyssee SA Licor infrared imaging system. For antibody detection membranes were washed with water and PBS and blocked in PBS with 5% milk powder. Immunoblots were probed concomitantly with affinity purified anti PGL-1, anti PGL-3 (clone E81 and clone F40, 0.2 *µg/ml*, MPI-CBG antibody facility, Dresden) and anti tubulin (clone DM1A, 1:20000, Sigmaaldrich) mouse monoclonal antibodies in PBST with 5% milk powder overnight at 4°C, washed five times, probed with secondary anti mouse antibody (IRdye 800CW, 1:20000, Licor) for 1h, washed five times with PBST, rinsed in PBS and imaged at 800nm in an Odyssee SA Licor infrared imaging system. Quantification of protein bands from Odyssee images (representative image see Fig. S3 B) was done with the Gel Analyzer tool in Fiji[**?**] (Rasband, W.S., ImageJ, U. S. National Institutes of Health, Bethesda, Maryland, USA, https://imagej.nih.gov/ij/, 1997-2018). Full lanes were selected, plotted and the specific protein bands identified and measured from the plot. Values were normalized to the protein amount loaded in each lane, which was determined for the identical lanes by the revert stain as well as by the tubulin levels from the blot. Quantifications of PGL-1 and PGL-3 for each worm line were made by measuring three extract dilutions (corresponding to approx. 200, 400 and 600 embryos per lane) and each dilution value was normalized to the identical lanes by revert and tubulin and averaging the data. PGL-1 and PGL-3 protein concentrations were calculated by normalizing the values from five independent experiments to PGL-1 and PGL-3 concentrations determined by quantitative label free mass spectrometry from N2 wildtype early embryos [32].

### S5. BROOD SIZE ANALYSIS OF WORM STRAINS

The number of offspring for individual worms were measured for animals reared at 10, 13, 16, 19, 22, 25, 26, 27 and 28°C for each strains used in this study. Carefully staged L4 worms were picked from stock plates to single 6 cm NGM plates inoculated with OP50. This was repeated for 12 repeats for each worm strain and for each temperature tested. The plates were then placed in IPP30plus incubators (Memmert) set to the test temperature. Incubators were calibrated against a certified mercury thermometer. Periodically, the founder worms were picked on to new plates leaving recently laid eggs on the plate. All plates were returned to the test temperature and the process repeated until the animals stopped laying eggs. Once the first generation of offspring had reached the L3 or L4 stages, the number of worms were estimated using the WormScan software [**?**] and a Perfection V750 Pro scanner (Epson). Counts were made per plate then added together to give the total number of offsping per animal. Worms which crawled onto the side of the plate or were killed during the transfer process were excluded from the data set.

### S6. FERTILITY ASSAY

Carefully staged L4 worms were singled to NGM plates (see SI section S5). Plates were shifted to the test temperature and worms were allowed to lay eggs for 48 hours. The founder animals were picked off the plates leaving the first generation offspring on the plates. After a further three days of incubation at the test temperature, plates were visually inspected under the dissecting scope and 20 animals per condition were assigned to either of the two following categories: fertile, having a healthy-looking gonad or eggs in the uterus; or infertile having no gonad or no eggs in the uterus.

### S7. SI MOVIE S1

Temperature oscillations between 15 and 29 °C lead to dissolution and condensation of P granules labelled via PGL-1::GFP (corresponding to Fig. 2A in the main text). Presented data are 4 repeats and the shaded area represents the standard deviation. Furthermore, a homogenization of P granule sizes is visible over the course of the video with more small P granules condensing after bigger P granules have dissolved. Images are maximum projections of spinning-disc confocal images. Time in mm:ss and scale bar equals 5 *µm*.

**TABLE S1.** Fitting parameters for the piece-wise model

| | $A_1$ | $B_1$ [K] | $A_2$ | $B_2$ [K] | $\nu_1^0$ | $\nu_1^1$ [1/K] | $\nu_2^0$ | $\nu_2^1$ [1/K] |
| --- | --- | --- | --- | --- | --- | --- | --- | --- |
| WT | 0.484 | 6.15 | 0.484 | 6.15 | -288030 | 1232 | -288030 | 1232 |
| $\Delta pgl-3$ | 0.481 | 6.77 | 0.498 | 1.59 | -251220 | 1114 | -95722 | 761 |

**TABLE S2.** Fitting parameters obtained from global minimisation

| | $A$ | $B$ [K] | $\Delta A$ | $\Delta B$ [K] | $\nu_1$ | $\nu_2$ [1/K] | $\Delta \nu_1$ | $\Delta \nu_2$ [1/K] | $\theta_\chi$ | $\lambda_\chi$ | $\theta_\nu$ | $\lambda_\nu$ |
| --- | --- | --- | --- | --- | --- | --- | --- | --- | --- | --- | --- | --- |
| WT | 0.486 | 5.31 | 0 | 0 | -201702 | 931 | 0 | 0 | N/A | N/A | N/A | N/A |
| $\Delta pgl-3$ | 0.486 | 5.05 | .001 | -3.10 | -261877 | 1246 | 87385 | -214 | 291.1 | 1.87 | 291.5 | 1.98 |

**TABLE S3.** PGL-1-GFP phase diagram data (Fig. 3B)

| T [°C] | cin [$\mu M$] | cinSD [$\mu M$] | cout [$\mu M$] | coutSD [$\mu M$] | # Embryos with droplets | # Embryos | # Droplets total |
| --- | --- | --- | --- | --- | --- | --- | --- |
| 5 | 13.20 | 5.68 | 0.37 | 0.05 | 13 | 13 | 813 |
| 10 | 12.55 | 4.44 | 0.38 | 0.05 | 24 | 24 | 2255 |
| 15 | 10.64 | 5.22 | 0.40 | 0.07 | 21 | 21 | 1835 |
| 20 | 8.06 | 3.43 | 0.43 | 0.06 | 19 | 19 | 1275 |
| 25 | 7.07 | 3.14 | 0.46 | 0.07 | 18 | 19 | 394 |
| 31 | 5.26 | 1.66 | 0.46 | 0.10 | 9 | 32 | 26 |

**TABLE S4.** PGL-3-GFP Δpgl-1 phase diagram data (Fig. 3C)

| T [°C] | cin [$\mu M$] | cinSD [$\mu M$] | cout [$\mu M$] | coutSD [$\mu M$] | # Embryos with droplets | # Embryos | # Droplets total |
| --- | --- | --- | --- | --- | --- | --- | --- |
| 5 | 12.88 | 10.01 | 0.40 | 0.07 | 7 | 7 | 323 |
| 10 | 10.15 | 9.04 | 0.45 | 0.12 | 12 | 12 | 264 |
| 15 | 8.78 | 6.78 | 0.46 | 0.07 | 10 | 11 | 158 |
| 20 | 2.55 | 1.42 | 0.47 | 0.12 | 12 | 14 | 169 |
| 25 | 2.04 | 0.51 | 0.48 | 0.15 | 9 | 10 | 148 |
| 31 | 2.06 | 0.37 | 0.52 | 0.13 | 10 | 18 | 79 |

**FIG. S1.**
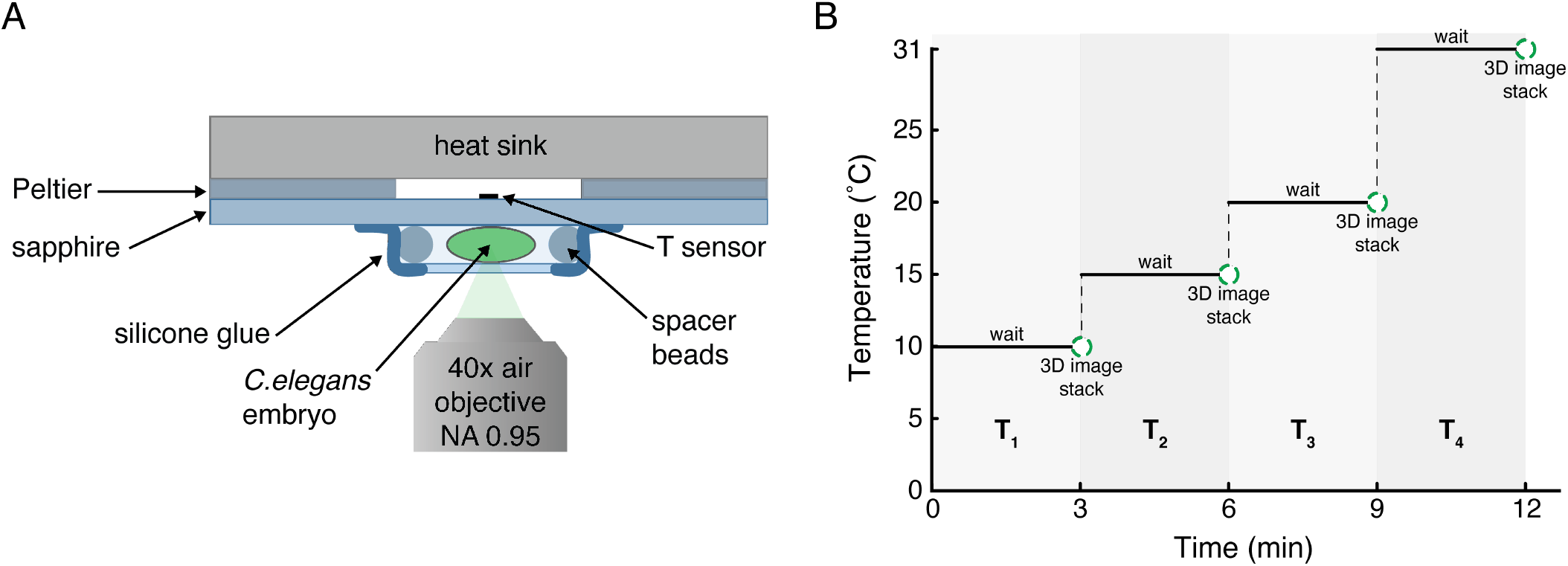
Temperature stage, mounting and experimental protocol. A. Feedback controlled global temperature stage capable of fast and precise temperature changes. Setup is based on a Sapphire glass connected to two Peltier modules that are computer controlled via a PID device connected to the T sensor. The Sapphire is scratch robust and quickly transmits the temperature of the Peltier to a transparent surface. *C. elegans* embryos are mounted in a sandwich with a controlled thickness of 25 *µm* via polystytene spacer beads. Using an air objective ensures there is very little temperature change introduced by the temperature difference between objective and the sample, due to the low conductivity of air. B. Experimental procedure where the first temperatures *T*_1_, *T*_2_, *T*_3_ are chosen from a range of 5 to 25 °C in steps of 5 °C and the final *T*_4_ = 31 °C. Temperatures are chosen in a random fashion to exclude effects of the temperature sequence and timing in the developmental processes of an early one-cell embryo. Confocal image stack are taken after a waiting period of 3 minutes at the current temperature to assure local thermodynamic equilibrium (see also SI Fig. S7.

**FIG. S2.**
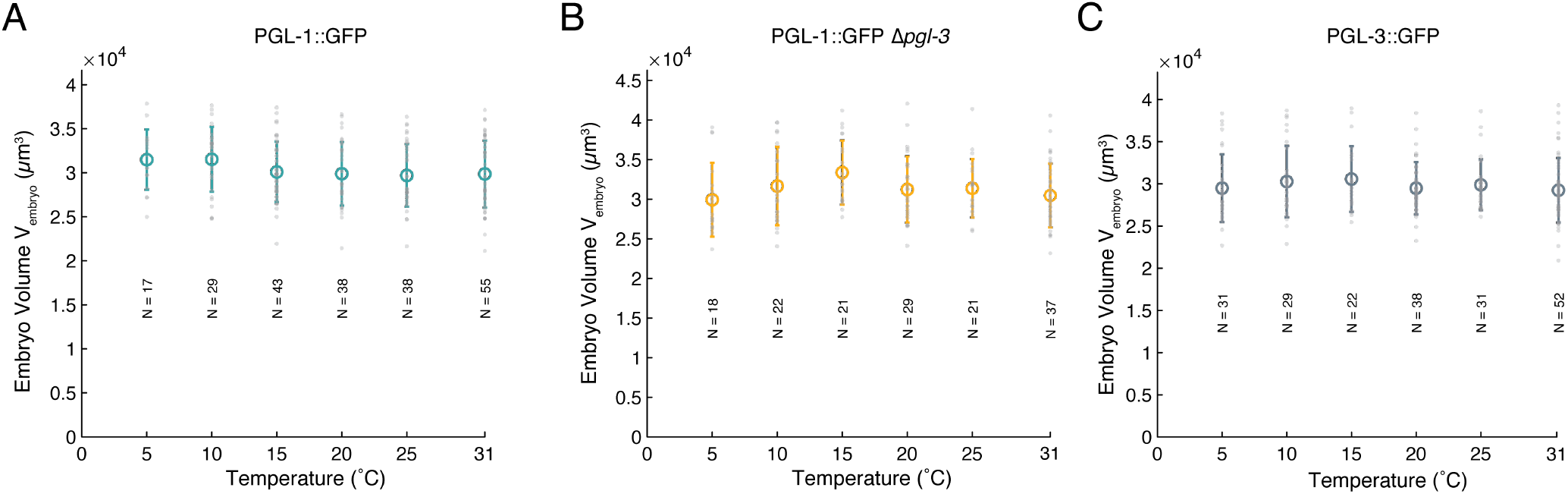
Temperature dependence of the one-cell embryo volume. For all worm strains used the total volume of an embryo is in a similar range of 30000 *µm*^3^. There is no significant change of the embryo volume in the temperature range that was used in this study. Error bars are standard deviation and N is the number of individual embryo volumes that were used for each temperature.

**FIG. S3.**
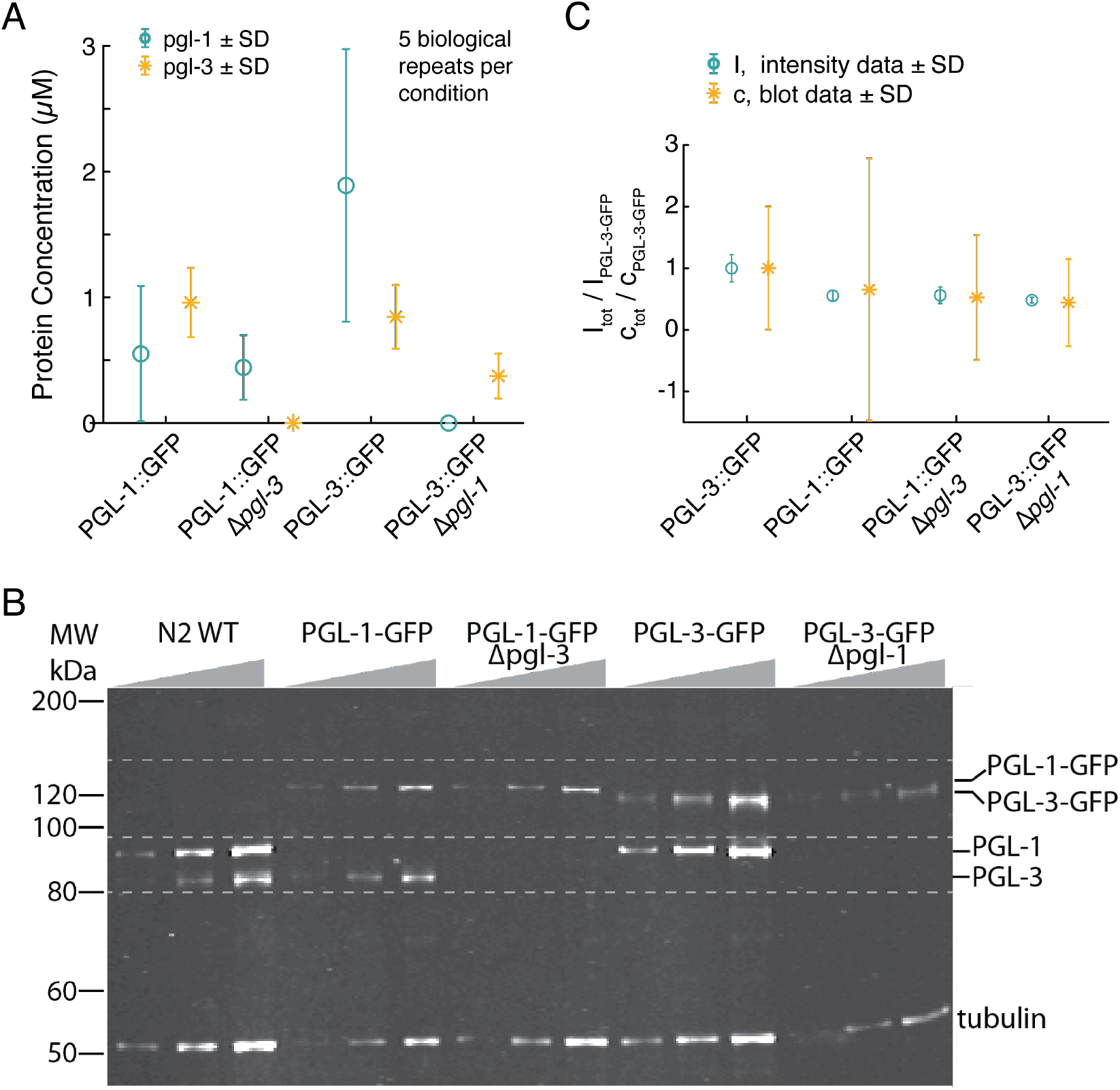
Average concentrations of PGL-1/3 of worm strains via western blots and average total intensity measurements. A.Data of average PGL-1 and PGL-3 concentrations via western blots from 5 biological repeats consisting of thousands of early embryos each (Error bars are SD). While PGL-3 concentration seems to be more conserved, PGL-1 is subjected to larger fluctuations between strains and a larger spread from biological repeats. B. Representative immunoblot from which protein concentrations were measured. Values were normalized and related to known concentrations from N2 worms [32]. C. Relative total embryo intensity *I* _total_ in comparison to the respective relative concentration as determined in A. Intensity values are relative to the PGL-3::GFP *I* _total_. Average total intensity values follow the same trend for the different strains as the corresponding western blot derived relative concentration values. However, *I* _total_ data has a reduced variability.

**FIG. S4.**
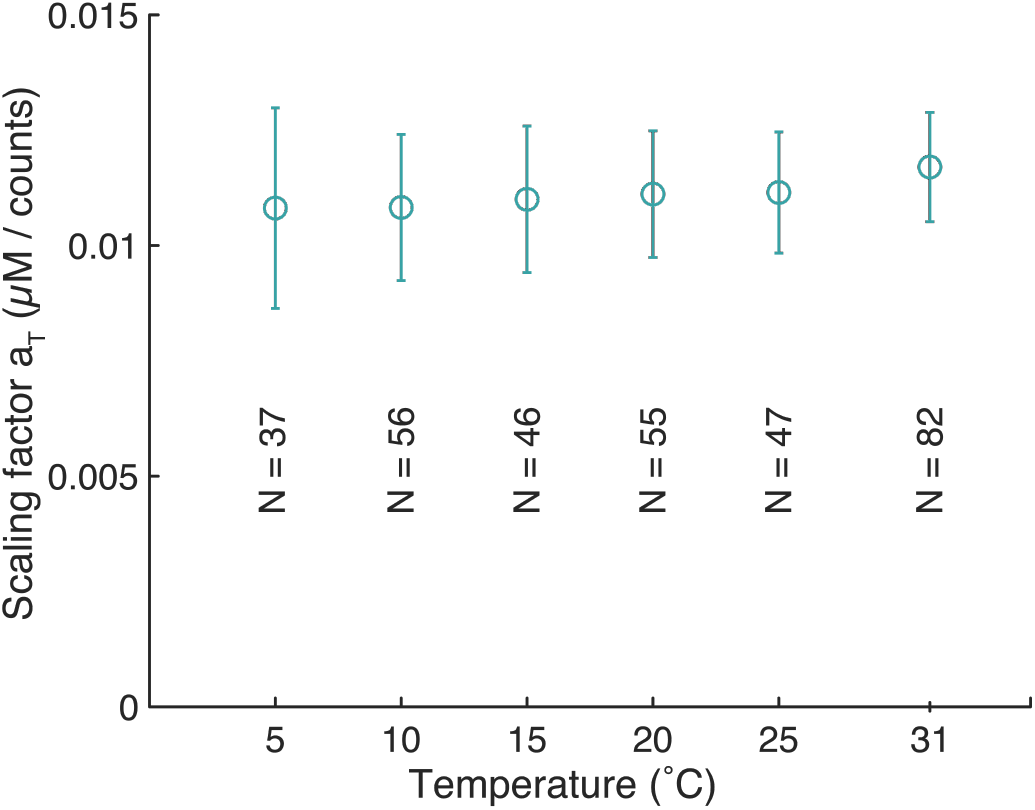
Scaling factor *aT* to relate fluorescence intensities with measured PGL-1 and PGL-3 protein concentrations. To calculate the relation all measured early one-cell embryos from all used worm strains were used and compared to the concentrations measured via immunoblotting (see Fig. SI S3). The influence of temperature on the scaling factor is not significant but as expected corresponds to slightly lower fluorescence intensities for higher temperatures.

**FIG. S5.**
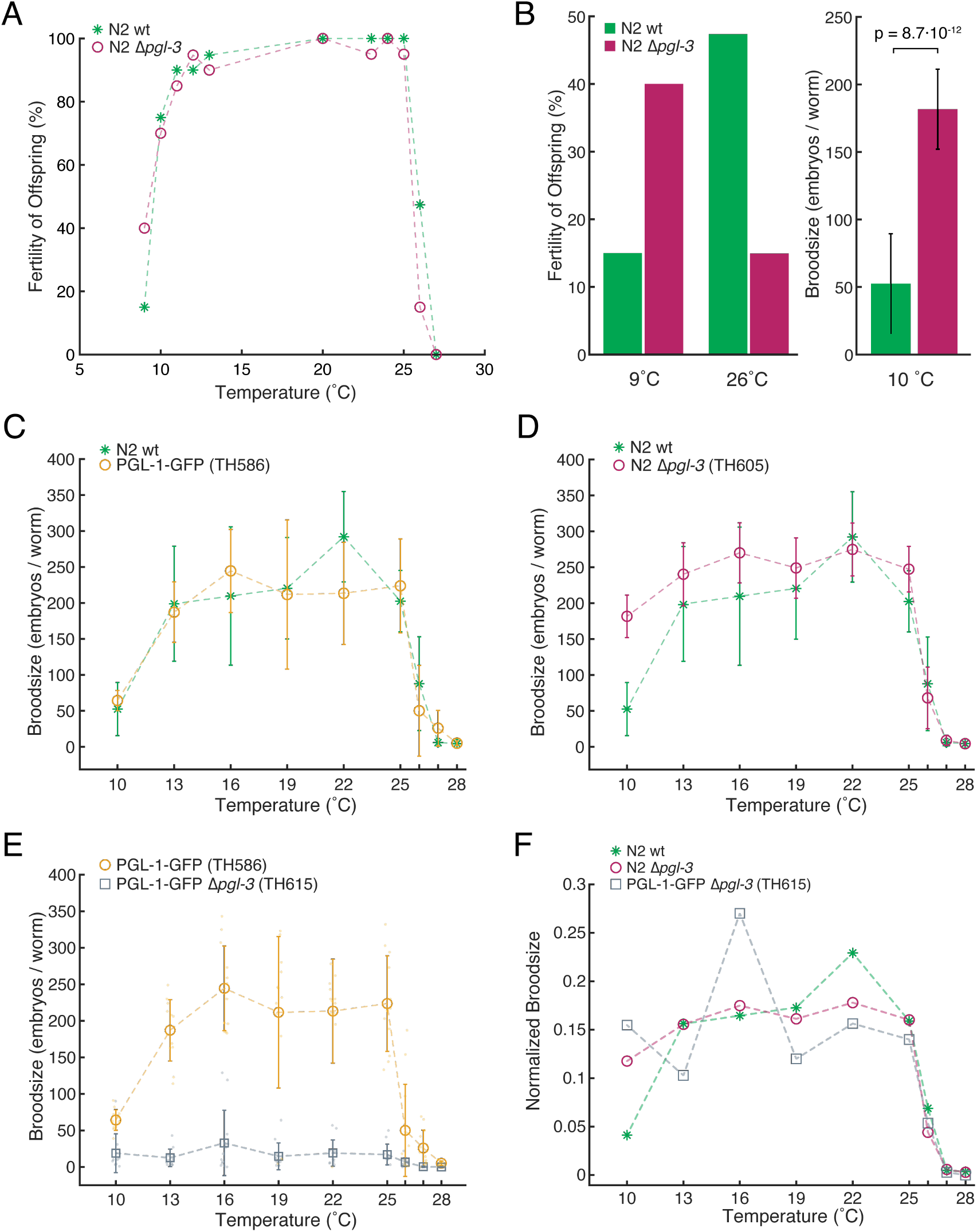
Temperature dependant broodsize and fertility. A. The fertility is a binary measure (gonad, no gonad) for the offspring of adult worms plated at specific temperatures. We see a similar fertility at moderate temperatures for the N2 ∆*pgl-3* strain (red circles) compared to wildtype N2 (green stars). Similar to the broodsize data in D we see an increase of fertility at low temperatures (at 9 °C) however additionally a decrease of fertility at 26 °C (20 experiments for each data point, also see B). B. Comparison of selected data from panel A and D, which indicates differences for N2 ∆*pgl-3* (red) versus N2 (green) at low and high temperature flanks (p-values are given for a two-sample t-test). C. PGL-1-GFP strain (yellow circles) and wildtype N2 (green stars) broodsize show equal behaviour in the tested temperature range (12 repeats each, errorbars are standard deviation). D. N2 ∆*pgl-3* (red cicles) and wildtype N2 broodsize show similar behaviour over most of the temperature range, while at 10 °C the ∆*pgl-3* strain has a significantly higher broodsize (12 repeats each, errorbars are standard deviation, also see B). E. Broodsize for strains used in the main document where PGL-1-GFP ∆*pgl-3* shows a strong decrease in brood size compared to the PGl-1-GFP (grey squares) strain (12 repeats each, errorbars are standard deviation). F. When normalizing the broodsize with the sum of embryos over all temperatures for each strain we see a similar trend for the N2 ∆*pgl-3* and PGL-1-GFP ∆*pgl-3* strain.

**FIG. S6.**
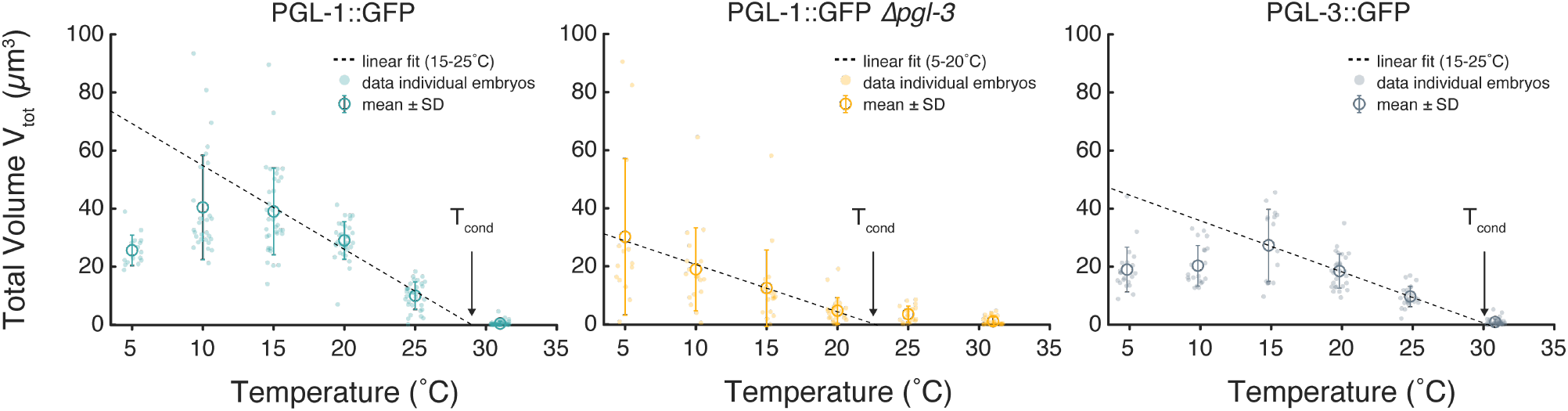
Total P granule volume *V*_*tot*_ versus temperature. For each temperature the mean and standard deviation as well as individual embryo data are presented. The linear regression to the individual data points is adopted to the approximate linear region as indicated in the legend of each strain. While for the Δ*pgl-3* deletion strain there seems to be a constant increase in total volume from 20 to 5 °C the PGL-1-GFP and PGL-3-GFP strain seem to flatten or decrease the total volume from 10 °C downwards. The intersect with the *V*_*tot*_ = 0 line represents the average condensation temperature for the whole ensemble of each worm strain.

**FIG. S7.**
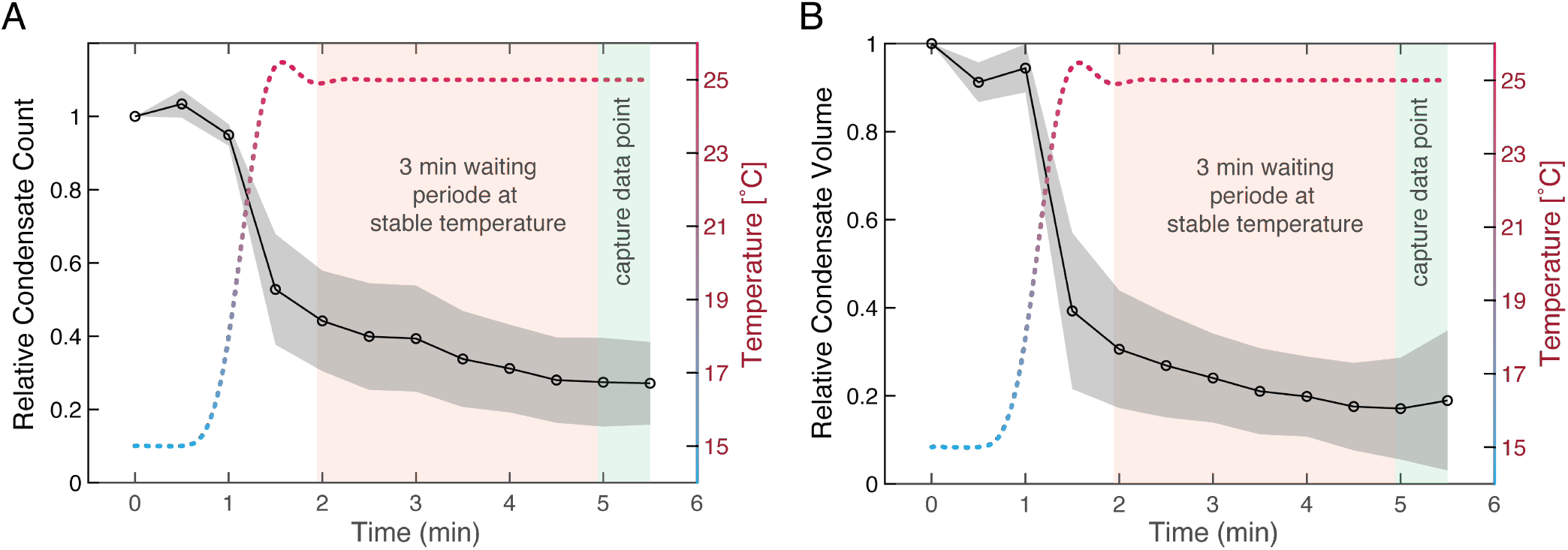
Control measurements for the protocol (see S1) used to derive volume and phase diagram data throughout this manuscript. Data shown is comprised of 6 embryo repeats with standard deviation and is normalised to the first data point. Temperatures are measured via a sensor of the temperature stage in sync with the image acquisition. A. Relative amount of P granule condensates as response to a temperature shock from 15 to 25 °C. Red shaded area depicts the waiting period of 3 min after the temperature settled. After this waiting period data was collected in the measurements that are presented in the main manuscript. B. Relative amount of P granule volume as response to a temperature shock from 15 to 25 °C.

**FIG. S8.**
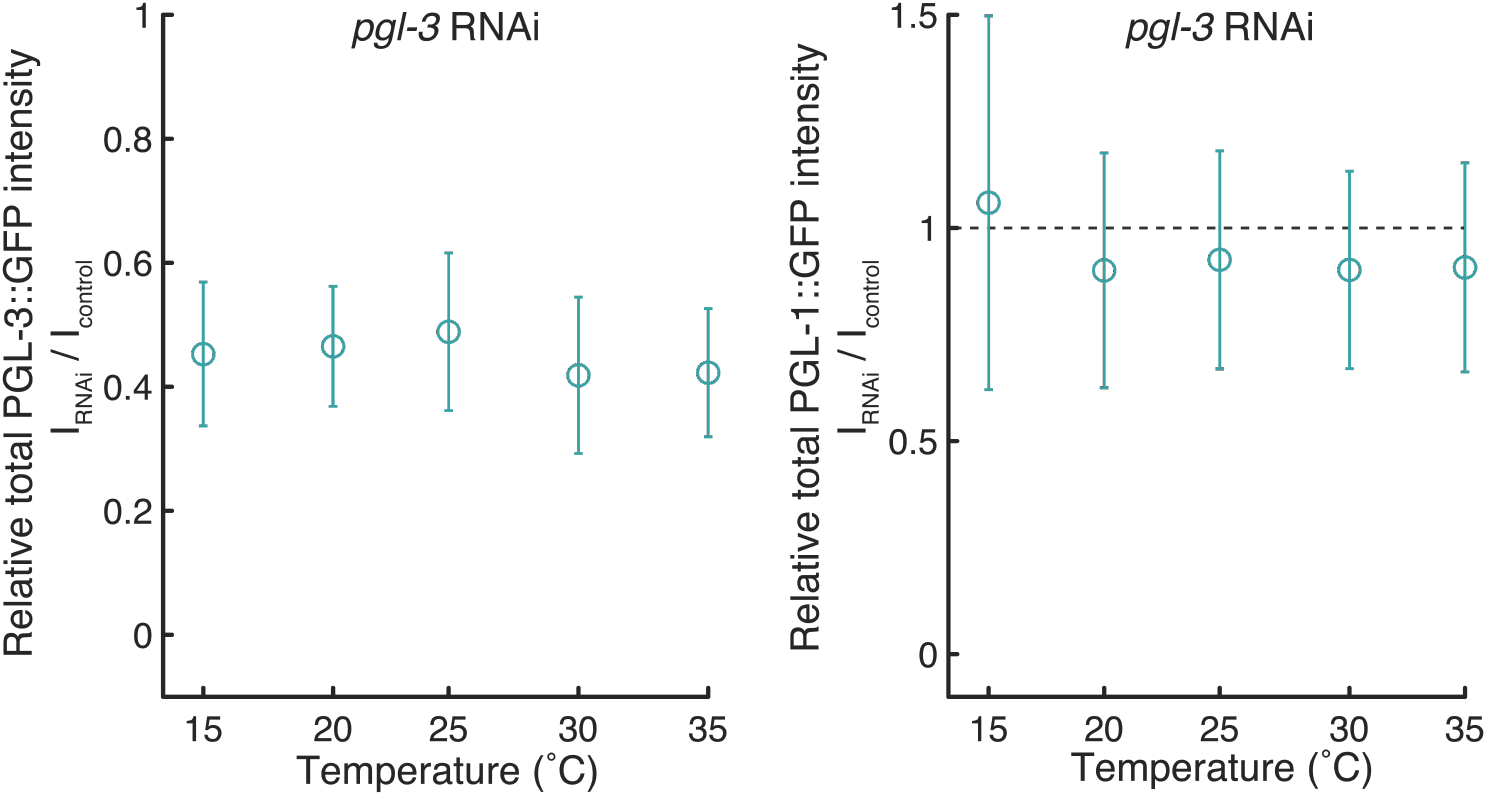
To quantify the amount of reduction in PGL-3 induced via *pgl-3* RNAi both PGL-1::GFP and PGL-3::GFP worm strains were subjected to the same RNAi protocol at the same time. A. We then quantified the RNAi relative effect on the total intensities *I*_*total*_ in the RNAi treated PGL-3::GFP strain against an untreated control. A significant decrease in relative total intensities for PGL-3::GFP under the influence of the RNAI is visible. B. Similar to A we also quantified the effect of *pgl-3* RNAi on PGL-1::GFP compared to an untreated control. As expected, the amount of PGL-1::GFP is not significantly affected the *pgl-3* RNAi treatment. In both cases there is no significant influence of short term temperature changes on the RNAi effect. Data points show average values and standard deviation versus temperature.

## References

[1] A. A. Hyman, C. A. Weber, and F. Jülicher, Annual review of cell and developmental biology 30, 39 (2014).

[2] S. Banjade and M. K. Rosen, eLife 3 (2014).

[3] S. F. Banani, H. O. Lee, A. A. Hyman, and M. K. Rosen, Nature reviews Molecular cell biology 18, 285 (2017).

[4] D. Dormann, The Journal of Cell Biology 219 (2019), ISSN 0021-9525, e201910211, https://rupress.org/jcb/article-pdf/219/1/e201910211/841178/jcb201910211.pdf, URL https://doi.org/10.1083/jcb.201910211.

[5] M. G.-J. Navarro, S. Kashida, and R. e. a. Chouaib, Nature Communications 10 (2019).

[6] C. P. Brangwynne, C. R. Eckmann, D. S. Courson, A. Rybarska, C. Hoege, J. Gharakhani, F. Jülicher, and A. A. Hyman, Science 324, 1729 (2009), ISSN 00368075.

[7] D. Updike and S. Strome, in Journal of Andrology (2010), vol. 31, pp. 53–60, ISSN 01963635.

[8] M. Hondele, R. Sachdev, S. Heinrich, J. Wang, P. Vallotton, B. M. Fontoura, and K. Weis, Nature 573, 144 (2019).

[9] J. Howard, Mechanics of Motor Proteins and the Cytoskeleton (Sinauer Associates, Publishers, 2001), ISBN 978-0-87893-334-1.

[10] F. Jülicher, A. Ajdari, and J. Prost, Reviews of Modern Physics 69, 1269 (1997).

[11] C. A. Weber, D. Zwicker, F. Jülicher, and C. F. Lee, Reports on Progress in Physics 82, 064601 (2019).

[12] A. F. D. Delgadillo, PhD thesis at TU Dresden none (2016).

[13] A. Putnam, M. Cassani, J. Smith, and G. Seydoux, Nature structural & molecular biology 26, 220 (2019).

[14] M. L. Begasse, M. Leaver, F. Vazquez, S. W. Grill, and A. A. Hyman, Cell reports 10, 647 (2015), ISSN 2211-1247.

[15] M. L. Huggins, The Journal of Physical Chemistry 9, 441 (1941).

[16] P. J. Flory, The Journal of chemical physics 10, 51 (1942).

[17] M. Rubinstein, R. H. Colby, et al., Polymer physics, vol. 23 (Oxford university press New York, 2003).

[18] I. Kawasaki, A. Amiri, Y. Fan, N. Meyer, S. Dunkelbarger, T. Motohashi, T. Karashima, O. Bossinger, and S. Strome, Genetics 167, 645 (2004), ISSN 0016-6731, 1943-2631, publisher: Genetics Section: Investigations, URL https://www.genetics.org/content/167/2/645.

[19] What is the power consumption of a cell?, http://book.bionumbers.org/what-is-the-power-consumption-of-a-cell/ (2020), accessed: 2020-10-22.

[20] A. M. Makarieva, V. G. Gorshkov, B.-L. Li, S. L. Chown, P. B. Reich, and V. M. Gavrilov, Proceedings of the National Academy of Sciences 105, 16994 (2008).

[21] A. Thommen, S. Werner, O. Frank, J. Philipp, O. Knittelfelder, Y. Quek, K. Fahmy, A. Shevchenko, B. M. Friedrich, F. Jülicher, et al., Elife 8, e38187 (2019).

[22] J. Blumm and A. Lindemann, High Temp. High Press 35, 627 (2003).

[23] M. E. Peterson, R. M. Daniel, M. J. Danson, and R. Eisenthal, Biochemical Journal 402, 331 (2007).

[24] D. Applegate, Journal of Muscle Research & Cell Motility 10, 457 (1989).

[25] S. B. Rivera, S. J. Koch, J. M. Bauer, J. M. Edwards, and G. D. Bachand, Fungal Genetics and Biology 44, 1170 (2007).

[26] J. Prost, F. Jülicher, and J.-F. Joanny, Nature physics 11, 111 (2015).

[27] F. Jülicher, S. W. Grill, and G. Salbreux, Reports on Progress in Physics 81, 076601 (2018).

[28] M. Mittasch, P. Gross, M. Nestler, A. W. Fritsch, C. Iserman, M. Kar, M. Munder, A. Voigt, S. Alberti, S. W. Grill, et al., Nature Cell Biology 20, 344 (2018), ISSN 1476-4679, URL https://www.nature.com/articles/s41556-017-0032-9.

[29] R. S. Kamath, A. G. Fraser, Y. Dong, G. Poulin, R. Durbin, M. Gotta, A. Kanapin, N. Le Bot, S. Moreno, M. Sohrmann, et al., Nature 421, 231 (2003), ISSN 1476-4687, number: 6920 Publisher: Nature Publishing Group, URL https://www.nature.com/articles/nature01278.

[30] H. Min, Y.-H. Shim, and I. Kawasaki, Journal of Cell Science 129, 341 (2016), ISSN 0021-9533, https://jcs.biologists.org/content/129/2/341.full.pdf, URL https://jcs.biologists.org/content/129/2/341.

[31] J. A. Arribere, R. T. Bell, B. X. H. Fu, K. L. Artiles, P. S. Hartman, and A. Z. Fire, Genetics 198, 837 (2014), ISSN 0016-6731, 1943-2631, publisher: Genetics Section: Investigations, URL https://www.genetics.org/content/198/3/837.

[32] S. Saha, C. A. Weber, M. Nousch, O. Adame-Arana, C. Hoege, M. Y. Hein, E. Osborne-Nishimura, J. Mahamid, M. Jahnel, L. Jawerth, et al., Cell 166, 1572 (2016), ISSN 10974172.

[33] S. Safran, Statistical thermodynamics of surfaces, interfaces, and membranes (CRC Press, 2018).

[34] U. Cho, S. M. Zimmerman, L.-c. Chen, E. Owen, J. V. Kim, S. K. Kim, and T. J. Wandless, PLoS One 8, e72393 (2013).

[35] E. E. Griffin, D. J. Odde, and G. Seydoux, Cell 146, 955 (2011).

